# ROR2 regulates cellular plasticity in pancreatic neoplasia and adenocarcinoma

**DOI:** 10.1101/2023.12.13.571566

**Authors:** Simone Benitz, Alec Steep, Malak Nasser, Jonathan Preall, Ujjwal M. Mahajan, Holly McQuithey, Ian Loveless, Erick T. Davis, Hui-Ju Wen, Daniel W. Long, Thomas Metzler, Samuel Zwernik, Michaela Louw, Donald Rempinski, Daniel Salas-Escabillas, Sydney Brender, Linghao Song, Ling Huang, Zhenyu Zhang, Nina G. Steele, Ivonne Regel, Filip Bednar, Howard C. Crawford

## Abstract

Cellular plasticity is a hallmark of pancreatic ductal adenocarcinoma (PDAC) starting from the conversion of normal cells into precancerous lesions to the progression of carcinoma subtypes associated with aggressiveness and therapeutic response. We discovered that normal acinar cell differentiation, maintained by the transcription factor Pdx1, suppresses a broad gastric cell identity that is maintained in metaplasia, neoplasia, and the classical subtype of PDAC in mouse and human. We have identified the receptor tyrosine kinase Ror2 as marker of a gastric metaplasia (SPEM)-like identity in the pancreas. Ablation of *Ror2* in a mouse model of pancreatic tumorigenesis promoted a switch to a gastric pit cell identity that largely persisted through progression to the classical subtype of PDAC. In both human and mouse pancreatic cancer, ROR2 activity continued to antagonize the gastric pit cell identity, strongly promoting an epithelial to mesenchymal transition, conferring resistance to KRAS inhibition, and vulnerability to AKT inhibition.

**Significance:** We discovered the receptor tyrosine kinase ROR2 as an important regulator of cellular identity in pancreatic precancerous lesions and pancreatic cancer. ROR2 drives an aggressive PDAC phenotype and confers resistance to Kras inhibitors, suggesting that targeting ROR2 will enhance sensitivity to this new generation of targeted therapies.

## INTRODUCTION

Pancreatic ductal adenocarcinoma (PDAC) is a deadly disease with few curative options for the vast majority of patients, in part because early detection methods are lacking. PDAC is marked by extensive tumor cell heterogeneity comprised of “classical” and “basal-like” molecular subtypes that, instead of acting as stable cellular communities, exist on a spectrum, with plasticity between subtypes likely contributing to therapeutic resistance [1–3]. Cellular plasticity is a hallmark of pancreatic cancer throughout progression, including transformation of normal cells into precancerous metaplastic lesions accompanied by the acquisition of distinct cellular identities. Whether cellular plasticity in precancerous disease is reflected in late-stage cancer progression has not been thoroughly considered, though some studies suggest a connection [4, 5].

Both pancreatic duct and acinar cells can give rise to PDAC in mouse models [6–8]. Acinar cells are more amenable to transformation in mice [8] and epigenetic analysis in patient samples suggests an acinar cell of origin [9]. Inflammation or expression of oncogenic Kras^G12D^ induces transdifferentiation of acinar cells to duct-like progenitor cells, a process called acinar-to-ductal metaplasia (ADM) [6, 7, 10]. Metaplastic duct cells are characterized by an up-regulation of genes that define normal duct cells, such as *Krt19* and *Sox9* and by loss of acinar differentiation markers, such as those encoding digestive enzymes [8, 11, 12].

The great majority of adult acinar cells are resistant to Kras^G12D^-induced transformation, but in combination with the ablation of transcription factors that otherwise maintain acinar cell identity, such as *Hnf1a*, *Nr5a2*, *Mist1*, *Ptf1a* and *Pdx1*, transformation is significantly accelerated [13–17]. Of these, genetic variants in *HNF1A*, *NR5A2* and *PDX1* are associated with susceptibility to PDAC development in patients [15, 18]. These findings support the hypothesis that maintenance of acinar cell identity is a powerful suppressor of pancreatic cancer initiation.

In both injury and neoplasia models, metaplastic ducts show a high degree of cellular heterogeneity marked by a variety of gastric cell identities including gastric chief-, pit-, tuft-, and enteroendocrine-like cells, together with subpopulations of senescent and proliferative metaplastic epithelia [19]. In experimental pancreatitis, a subset of meta/dysplastic cells take on a so-called SPEM (Spasmolytic Polypeptide-Expressing Metaplasia) identity [20], which is generally associated with gastric wound healing [21]. While gastric cell identity is evident in precancerous lesions, it has not been noted in carcinoma, which are instead marked by the classical and basal-like subtypes. The classical subtype exhibits higher expression of pancreatic lineage- and epithelial-defining transcription factors, whereas the basal-like subtype is characterized by an erosion of epithelial cell identity and acquisition of mesenchymal and basal cell features [22–26]. Importantly, subtype identity of pancreatic cancer cells is associated with prognosis, with the basal-like subtype correlating to poorer overall survival [1].

The complex heterogeneity in both precancerous lesions and carcinoma has been of intense interest. Unfortunately, the study of the earliest molecular changes associated with the reprogramming of normal pancreatic acinar cells to neoplasia has been challenging using standard single-cell RNA-sequencing methods, largely due to the fragility of normal acinar cells during tissue disruption, together with high levels of digestive enzymes, especially nucleases. To overcome this, we performed single-nucleus ATAC-sequencing of adult mouse pancreas tissue to identify possible mechanisms of cellular reprogramming initiated by the inducible ablation of the acinar cell identity factor *Pdx1*, the expression of oncogenic Kras^G12D^, or both together. Strikingly, chromatin accessibility of marker genes was sufficient to determine cell identity of multiple cell types within the tissues, including epithelial, fibroblast and immune cell populations. Interestingly, in transformation-sensitive *Pdx1*-null acinar cells, we identified increased accessibility and expression of several genes associated with a gastric SPEM signature, which was intensified in acinar cells also expressing *Kras^G12D^*. Among these genes, we focused on *Ror2*, which encodes a receptor tyrosine kinase in the noncanonical *Wnt* signal transduction pathway. Genetic ablation of *Ror2* in a mouse model of pancreatic neoplasia shifted cell identity, giving rise to precancerous lesions dominated by mucinous gastric pit-like cells. In PDAC, *ROR2* expression strongly correlated with a basal-like/mesenchymal phenotype where it controls cell proliferation. Forcing *ROR2* expression in classical subtype PDAC cell lines was sufficient to induce a complete conversion to rapidly proliferating, poorly differentiated mesenchymal cells. The concomitant shift in signal transduction induced by ROR2 expression conferred resistance to a KRAS^G12D^ inhibitor while introducing a profound sensitivity to AKT inhibition.

## RESULTS

### Tracking cell identity changes in early pancreatic cancer development using snATAC-seq

To initiate pancreatic tumorigenesis, oncogenic Kras must overcome resistance to transformation posed by acinar cell identity [13–17, 27]. In order to study the earliest steps in this process, we compared pancreata with conditional adult acinar cell specific expression of Kras^G12D^ alone to those also lacking Pdx1, a developmental transcription factor that we have shown previously to be important for maintaining acinar cell differentiation [13]. We used *Ptf1aCre^ERT^*-driven recombination to activate *Kras^G12D^* expression and ablate *Pdx1* by treating 8-to 9-week-old mice with Tamoxifen, followed by tissue collection 2 and 10 weeks later to capture early cell identity changes. Sole expression of *Cre^ERT^* (*Ptf1a^ERT^*) and loss of *Pdx1* (*Ptf1a^ERT^;Pdx1^f/f^*) did not cause any obvious morphological changes in the epithelial compartment, with the majority of cells being amylase-positive acinar cells (**Fig. 1A, 1B**). Oncogenic Kras expression in *Ptf1aCre^ERT^;Kras^LSL-G12D^* (*Ptf1a^ERT^;K\**) mice led to sporadic acinar-to-ductal metaplasia, characterized by loss of amylase, induced expression of the ductal marker Krt19, and a surrounding stromal reaction indicated by vimentin staining (**Fig. 1A, 1B**). In contrast to the low transformation efficiency of Kras^G12D^ alone, the concomitant loss of Pdx1 significantly accelerated cellular reprogramming in *Ptf1aCre^ERT^;Kras^LSL-G12D^;Pdx1^flox/flox^* (*Ptf1a^ERT^;K*;Pdx1^f/f^*) animals, as previously reported [13]. Only 10 weeks after Tamoxifen, acinar cells completely transdifferentiated into metaplastic duct-like cells, accompanied by a fibroinflammatory stromal reaction (**Fig. 1A, 1B**).

**Fig. 1:**
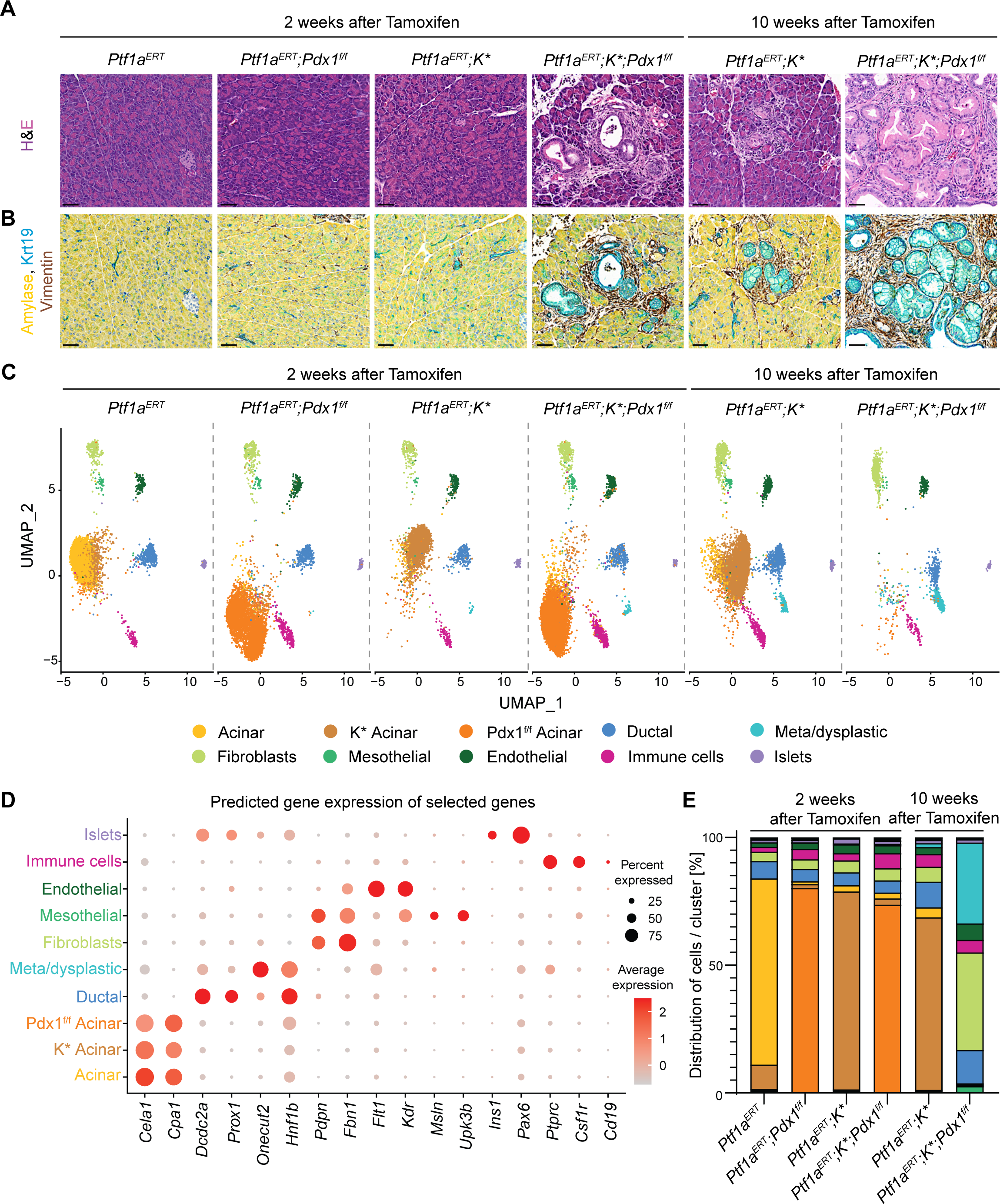
(**A**) Representative H&E staining of indicated mouse genotypes. Eight-to 9-week-old animals were given Tamoxifen to induce *Cre^ERT^*-mediated recombination and were sacrificed 2 and 10 weeks later. Scale bars, 50 µm. (**B**) Representative multiplex IHC staining for Amylase (yellow), Krt19 (teal) and Vimentin (brown). Scale bars, 50 µm. (**C**) Unsupervised clustering of snATAC-seq data from indicated mouse genotypes (n=2 each), represented as UMAP plots. Indicated cell types were identified. (**D**) Dot plot showing average gene expression and percentage of cells expressing selected marker genes across all identified clusters. (**E**) Percentage distribution of nuclei in each annotated cell cluster was determined over the total amount of nuclei in each genotype and time point (n=2).

To better define how Kras^G12D^ expression alone and in combination with Pdx1 ablation impacts acinar cell identity, we established a novel protocol allowing high-quality snATAC-seq of snap-frozen, rapidly pulverized bulk pancreatic tissue (**Suppl. Fig. 1A**). After extensive quality control and removal of low-quality nuclei (**Suppl. Fig. 1B, 1C**), we performed downstream analysis with a total of 33,505 nuclei and compared chromatin accessibility states of *Ptf1a^ERT^* and *Ptf1a^ERT^;Pdx1^f/f^* control animals to mice with *Kras^G12D^* expression, sacrificed 2 and 10 weeks post Tamoxifen.

Based on chromatin accessibility at the gene body and promoter, we calculated predicted gene expression activity scores by using the Signac pipeline [28] and used top predicted differentially accessible genes to assign cell cluster identity (**Suppl. Fig. 1D, 1E**). We identified acinar, ductal and meta/dysplastic cells as well as fibroblasts, mesothelial, endothelial, immune and islet cells (**Fig. 1C, Suppl. Fig. 1E**). Surprisingly, although appearing morphologically normal, acinar cells of different mouse genotypes grouped into very distinct cell clusters, while other cell types remained unperturbed across genotypes at this resolution. Notably, acinar cells of *Ptf1a^ERT^* control (Acinar, yellow cluster) and *Ptf1a^ERT^;K\** (K* Acinar, brown cluster) mice showed slight differences, while acinar cells of Pdx1 knockout mice (Pdx1^f/f^ Acinar, orange cluster) exhibited an obvious switch of cell identity, independent of Kras status (**Fig. 1C, Suppl. Fig. 1E**). Acinar cell clusters were defined by predicted expression of digestive enzymes, such as *Cela1* and *Cpa1,* which show heterogeneous expression levels across the clusters, an indicator of distinct differentiation states (**Fig. 1D**). Consistent with the histology, expression of Kras^G12D^ led to time- and genotype-dependent meta/dysplastic duct cell formation (teal cluster), characterized by expression of metaplastic and ductal marker genes, such as *Onecut2* and *Hnf1b*, respectively (**Fig. 1C, 1D, 1E**). These data were also consistent with the rampant transformation in *Ptf1a^ERT^;K*;Pdx1^f/f^* animals 10 weeks post Tamoxifen, with acinar cells being almost completely absent, replaced by an immense increase in meta/dysplastic duct cells (**Fig. 1C, 1E**). Of note, these massive changes in the epithelial compartment were accompanied by a substantial increase of fibroblasts (light-green cluster), with little change in immune cell numbers (violet cluster) (**Fig. 1E**).

Our snATAC-seq approach corroborates histological observations and changes in cell populations, that occur during early transformation of the pancreas. Importantly, we uncovered that acinar cells from different mouse genotypes segregate into distinct clusters with an obvious shift in identity in transformation-sensitive Pdx1-null acinar cells, revealing previously elusive details of early cellular reprogramming. We next sought to investigate gene accessibility changes among our identified acinar cell clusters in more detail.

### Ablation of Pdx1 initiates gastric transdifferentiation

To unmask cell identity differences across our acinar cell clusters, we subsetted acinar cells from the 2 week time point based on their genotype (**Fig. 2A**). To validate our identification of differentially regulated genes, we used an integrative approach combining our snATAC-seq with bulk RNA-seq data of isolated acinar cells from matching mouse genotypes and sex. Genes that showed the highest correlation across the two techniques are depicted in Figure 2B (**Fig. 2B, Suppl. Fig. 2A**). One gene confirming the validity of this approach was *Kdm6a*, which is located on the X chromosome and was both more accessible and more highly expressed in females (**Fig. 2B**). Acinar cells with Pdx1 loss exhibited an obvious decrease of accessibility and expression of acinar differentiation genes, such as *Cela1* or *Ptf1a* [13], but elevated levels of the AP-1 transcription factor *Junb*, which is associated with transcriptional reprogramming in ADM [29] (**Fig. 2B, Suppl. Fig. 2A**).

**Fig. 2:**
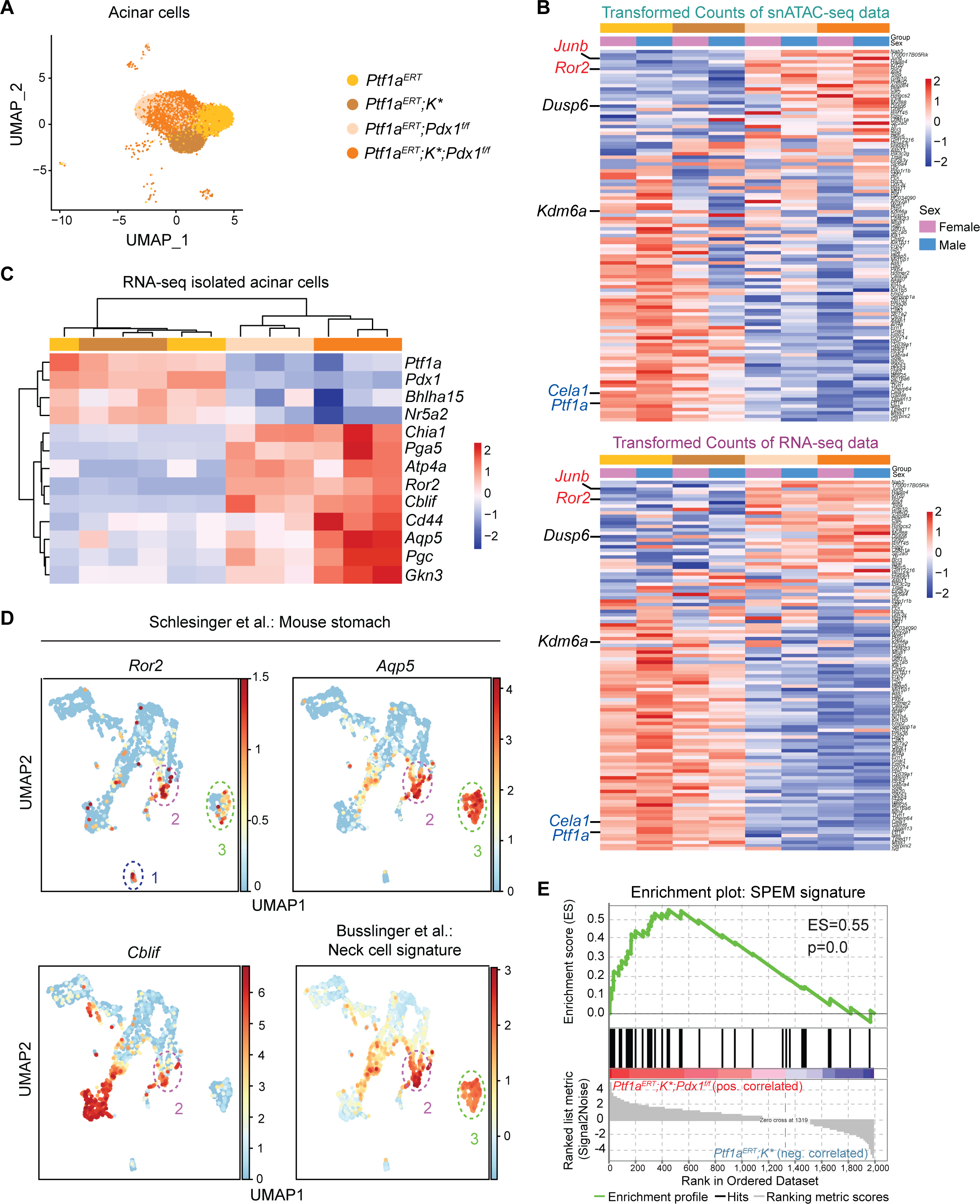
(**A**) UMAP analysis depicting acinar cell clusters from indicated mouse genotypes, 2 weeks post Tamoxifen. (**B**) Heatmaps showing expression levels of genes, that showed highest correlation in both snATAC-seq (upper panel) and bulk RNA-seq datasets (lower panel). Expression levels are represented by color coding. (**C**) Heatmap depicting mRNA expression of selected acinar and gastric lineage genes. (**D**) UMAP plots using scRNA-seq data from mouse stomach cells (Schlesinger et al.) with gastric cell type signature annotation (Busslinger et al.). (**E**) Gene set enrichment analysis results for SPEM signature comparing *Ptf1a^ERT^;K\** and *Ptf1a^ERT^;K*;Pdx1^f/f^* acinar cells.

Among the top genes that showed the most significant enrichment in accessibility and expression in transformation-sensitive *Pdx1*-null acinar cells was the receptor tyrosine kinase *Ror2* (Receptor Tyrosine Kinase Like Orphan Receptor 2) (**Fig. 2B, Suppl. Fig. 2A**). Particularly, Ror2 regulates non-canonical Wnt signaling and defines cancer cell identity in a variety of tumor types [30–32]. Besides *Ror2*, our RNA-seq data revealed that gastric lineage-specific genes, including *Cobalamin binding intrinsic factor* (*Cblif*), *Pepsinogen5* (*Pga5*), *Gastrokinin3* (*Gkn3*) and *Aquaporin5* (*Aqp5*) [33, 34] are up-regulated in *Pdx1*-knockout acinar cells (**Fig. 2C**). Considering that the acquisition of gastric signatures has recently been identified as a hallmark of pancreatic tumorigenesis [19, 35], we asked if Ror2 itself is associated with gastric cell identity.

Analysis of published mouse stomach scRNA-seq data [19] indeed revealed *Ror2* expression in gastric cell populations, which, in part, co-express *Cblif* and *Aqp5* and show enriched expression of enterochromaffin-like cell (ECL, population 1) and gastric neck cell signatures [33] (population 2 and 3) (**Fig. 2D, Suppl. Fig. 2B**). IHC staining confirmed Ror2 protein expression in the base and in Aqp5-positive neck cells of antral crypts in mouse stomach tissue (**Suppl. Fig. 2C**). Of note, gene expression patterns of deep antral cells share characteristics with Spasmolytic Polypeptide-Expressing Metaplasia (SPEM), which forms upon gastric injury as part of a wound-healing program, and that has been recently identified in pancreatitis-induced metaplasia [20]. In addition to *Ror2*, SPEM-associated genes, such as *Gkn3*, *Pgc* or *Aqp5*, were up-regulated in acinar cells with Pdx1 ablation, especially in combination with oncogenic Kras expression (**Fig. 2C**). Gene set enrichment analyses (GSEA) for previously published SPEM marker [20, 36–38] confirm that the SPEM signature is significantly up-regulated in *Ptf1a^ERT^;K*;Pdx1^f/f^* acinar cells (**Fig. 2E**). Thus, Pdx1 actively maintains acinar cell identity while it suppresses a gastric/SPEM identity and susceptibility to transformation by Kras. Notably, we identified *Ror2* as a previously-unrecognized marker of the gastric cell identity-associated gene signature.

### Ror2 marks gastric neck cell and SPEM signatures in pancreatic neoplasia

Given that oncogenic Kras-induced meta/dysplastic cells in the pancreas acquire features reminiscent of gastric cell lineages [19, 35], we examined if *Ror2* expression was associated with this identity shift. By performing RNA-ISH and IHC, we found that Ror2 is indeed expressed in a portion of meta/dysplastic cells in *Ptf1a^ERT^;K* animals*, comprising ∼25% of Krt19-positive cells, especially in later time points (**Fig. 3A**). To explore whether *Ror2* expression levels define meta/dysplastic duct cell identity, we used pancreas single-cell RNA-sequencing data from Schlesinger et al., which contained 3309 lineage-traced meta/dysplastic cells isolated from *Ptf1a^ERT^;K*;ROSA^LSL-TdTomato^* mice [19] (**Fig. 3B**). In concordance with our IHC data, *Ror2* is expressed in subpopulations of tdTomato-positive meta/dysplastic cells (**Fig. 3B**). To determine the identity of Ror2-positive cells, we analyzed for the expression of gastric cell type signatures from normal mouse gastric epithelia [33] as well as typical SPEM genes. We found that the Ror2^High^ cell population (**Fig. 3B**, circled) is indeed associated with higher expression of gastric neck cell and SPEM signatures. Consequently, we named this cluster neck/SPEM cell-like (**Fig. 3C**). Moreover, in accordance with Schlesinger et al. [19], we identified a pit cell-like population, confirmed by *Tff1* expression, and proliferating and senescent populations, exhibiting high expression of *Mki67* and *Cdkn2a*, respectively (**Fig. 3C, Suppl. Fig. 3A, 3B, 3C**). Three clusters did not exhibit clear segregation of marker gene expression, which we designate as “hybrid”. While the “acinar-like” cluster retained expression of acinar genes, expression of ductal genes was enriched in the “duct-like” cell cluster (data not shown). Of note, we found highest expression of *Ror2* in the neck/SPEM cell-like cluster, followed by dividing and acinar-like meta/dysplastic cell populations (**Fig. 3D**). Among the top 10 genes that showed the highest correlation to *Ror2* expression, we found the gastric neck and SPEM marker *Muc6* as well as the SPEM genes *Aqp5*, *Gkn3* or *Pgc* (**Fig. 3D, 3E**), reminiscent of the gene signature found in *Pdx1*-null acinar cells (**Fig. 2C, 2D**).

**Fig. 3:**
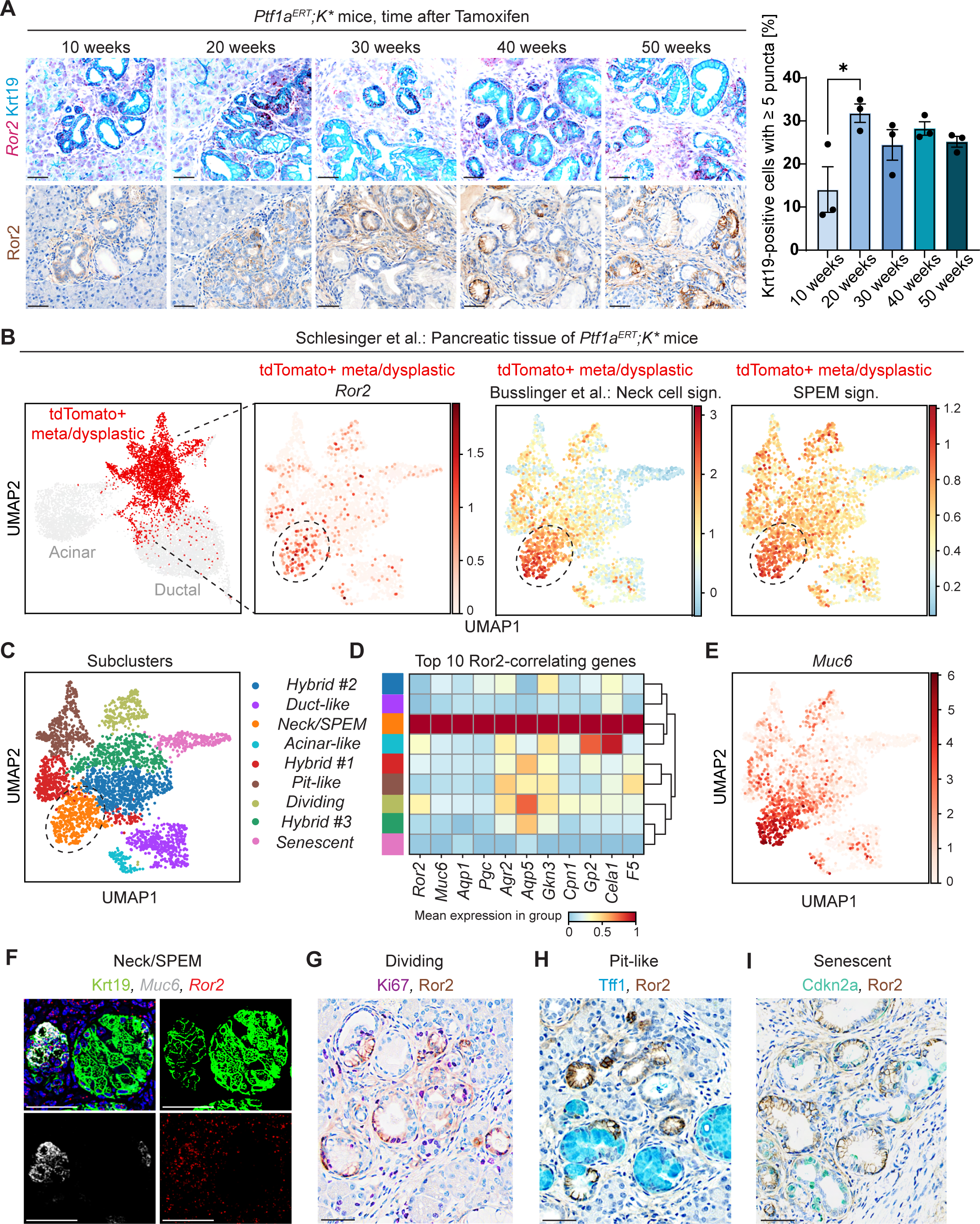
(**A**) Representative images of *Ror2* RNA-ISH in combination with Krt19 IHC (upper panel) and Ror2 IHC (lower panel) of pancreatic tissue from *Ptf1a^ERT^;K\** mice harvested at indicated time points post Tamoxifen. Scale bars, 50 μm. Percentage of Krt19-positive cells with ≥ 5 positive puncta was quantitated. All data are presented as mean ± SEM; p-values were calculated by two-tailed, unpaired Student’s *t*-test; * p < 0.05. (**B**) UMAP analysis depicting epithelial cell clusters of *Ptf1a^ERT^;K\** mice (left panel), Schlesinger et al. scRNA-seq data. TdTomato-positive meta/dysplastic cells were subsetted and *Ror2* expression and enrichment of gastric neck and SPEM cell signature is shown. (**C**) UMAP analysis illustrating subcluster identities of tdTomato-positive meta/dysplastic cells after integration with Harmony. (**D**) Heatmap representing expression of *Ror2* and of top 10 most significantly *Ror2*-correlating genes across identified Harmony clusters. (**E**) UMAP analysis showing expression of *Muc6*. (**F**) Multiplex IF and RNA-ISH staining for Krt19 (green), *Muc6* (white) and *Ror2* (red) on tissue of a *Ptf1a^ERT^;K\** mouse, 20 weeks post Tamoxifen. (**G**) Image showing Ki67 (purple) and Ror2 (brown) IHC. (**H**) Dual IHC depicting Tff1 (teal) and Ror2 (brown) staining. (**I**) Cdkn2a (green) and Ror2 (brown) dual IHC staining. For all staining, representative images are depicted. Scale bars, 50 µm.

Multiplex IF/ISH staining confirmed co-localization of *Ror2* with *Muc6* expression in ductal lesions of *Ptf1a^ERT^;K\** mice, 20 weeks post Tamoxifen (**Fig. 3F**). Moreover, we detected co-localization of Ror2 with Ki67 (**Fig. 3G**), showing that Ror2-positive meta/dysplastic cells are indeed associated with a proliferative neck/SPEM phenotype. In contrast, senescent and pit cell-like cells, identified by Cdkn2a and Tff1 IHC, respectively, were negative for Ror2 (**Fig. 3H, 3I**).

Interestingly, Ror2-positive lesions showed lower abundance of *Krt19* expression, consistent with the SPEM population found in injury-induced ADM [20] (**Fig. 3F, Suppl. Fig. 3D**). In the dataset from Ma et al. [20], a substantial proportion of “Mucin/Ductal” cells also expressed *Ror2*, again showing correlation with neck cell and SPEM marker gene expression, such as *Muc6* or *Aqp5* (**Suppl. Fig. 3E, 3F, 3G, 3H**). Overall, we discovered that Ror2 expression defines proliferative neck/SPEM cell-like subpopulations of pancreatic precancerous cells, while Ror2 negative cells exhibit senescent and pit cell-like characteristics. Since the relevance of these cellular identities on pancreatic cancer development is unknown, we next sought to determine if ablation of the neck/SPEM marker Ror2 impacts carcinogenesis in mice.

### Ror2 ablation shifts gastric metaplasia towards a pit cell identity

Next, to assess the functional relevance of Ror2 in early metaplasia and neoplasia, we generated *Ptf1a^ERT^;K\** mice with a conditional knockout allele of *Ror2* and a *ROSA^LSL-YFP^* reporter. *Ptf1aCre^ERT^*-mediated recombination led to the deletion of Exon3/4 of the *Ror2* gene in the majority of acinar cells (**Suppl. Fig. 4A**). IHC for the YFP reporter expression suggested a recombination efficiency of ∼90%. We noted that some lesions retained Ror2 expression, suggesting either less efficient recombination of this locus or possible selection for cells that escaped recombination (**Suppl. Fig. 4B**). While we observed slight differences in tissue remodeling, quantified as loss of acinar cells and formation of meta/dysplastic duct-like structures between *Ptf1a^ERT^;K\** and *Ptf1a^ERT^;K*,Ror2^f/f^* animals at 10 weeks post Tamoxifen (**Suppl. Fig. 4C)**, *Ptf1a^ERT^;K*,Ror2^f/f^* animals presented with significantly more transformation at 20 and 40 weeks post Tamoxifen (**Fig. 4A**). Notably, 40 weeks post Tamoxifen, 3 *Ptf1a^ERT^;K*,Ror2^f/f^* mice developed regions of carcinoma, a rapidity of progression not observed in our control cohort. They also showed a more pronounced stromal reaction with significantly elevated collagen deposition as represented by Picrosirius Red stain, which was also observed when *Ror2* was ablated in a mammary cancer model [39] (**Suppl. Fig. 4D**). When comparing lesion identity in mice that presented with neoplasia, we found that *Ptf1a^ERT^;K*,Ror2^f/f^* animals exhibit a striking increase in the number of mucinous meta/dysplastic lesions, the first indication of a possible shift in epithelial cell identity (**Fig. 4A**).

**Fig. 4:**
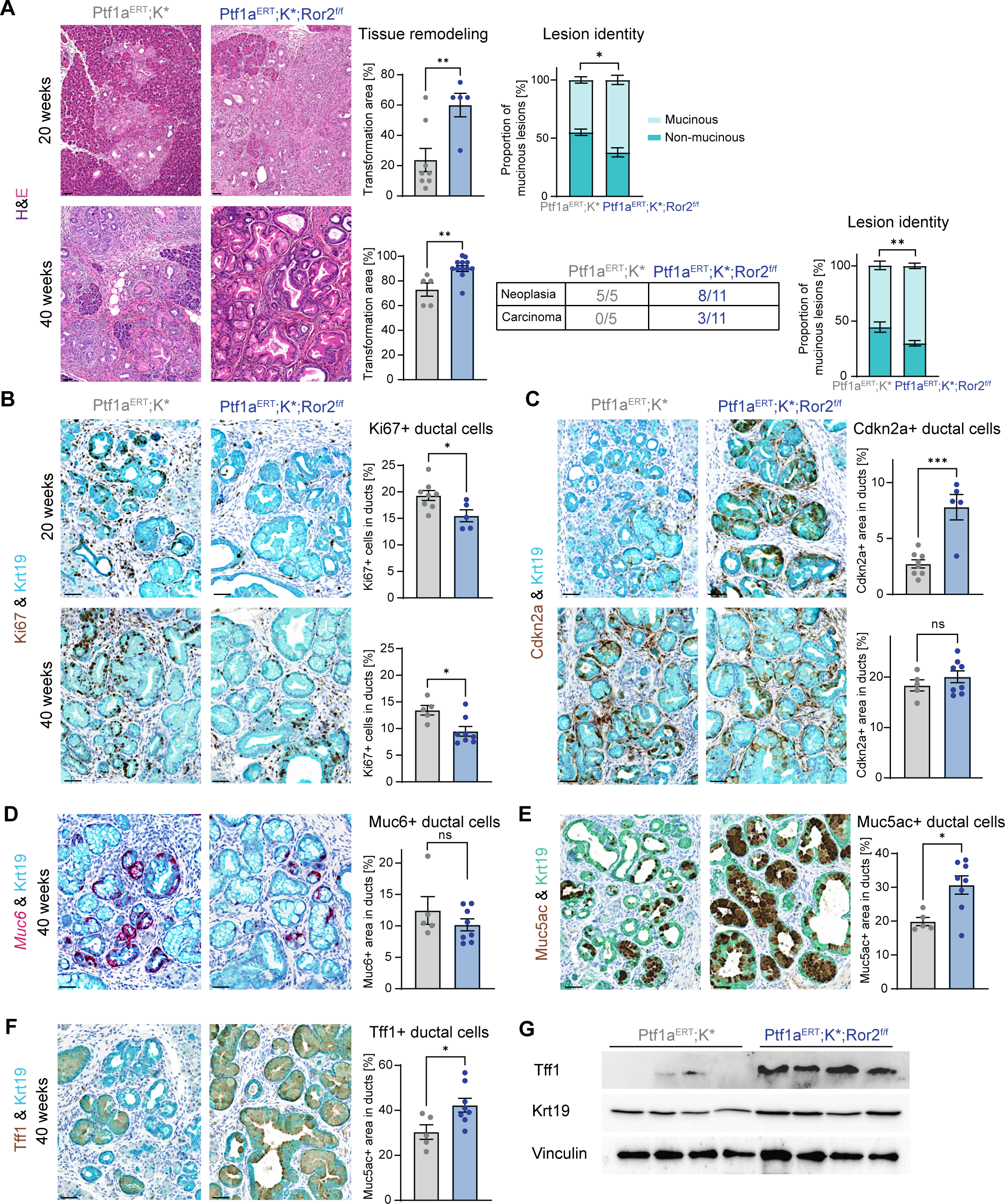
(**A**) H&E staining of *Ptf1a^ERT^;K\** and *Ptf1a^ERT^;K*;Ror2^f/f^* mice, 20 and 40 weeks after Tamoxifen administration. Tissue remodeling and number of mucinous and non-mucinous lesions were quantified (n=5-11). (**B**) Dual IHC staining for Ki67 (brown) and Krt19 (teal). (**C**) Images depicting dual Cdkn2a (brown) and Krt19 (teal) IHC staining. (**D**) Multiplex RNA-ISH and IHC staining detecting *Muc6* mRNA expression (red) and Krt19 (teal) in mice, sacrifized 40 weeks post Tamoxifen. (**E**) Dual IHC staining for Muc5ac (brown) and Krt19 (green). (**F**) Images showing Tff1 (brown) and Krt19 (teal) dual IHC. (**G**) Immunoblot analysis for Tff1 and Krt19 from protein lysates of *Ptf1a^ERT^;K\** and *Ptf1a^ERT^;K*;Ror2^f/f^* mice, sacrificed 40 weeks post Tamoxifen (n=4). Vinculin served as loading control. For all staining, representative images are depicted. Scale bars, 50 µm. Staining were quantified with HALO software and all data are presented as mean ± SEM; p-values were calculated by two-tailed, unpaired Student’s *t*-test; * p < 0.05, ** p < 0.01, *** p < 0.001.

Based on our previous observation that Ror2 was mainly expressed in proliferative neck/SPEM subpopulations of meta/dysplastic cells, but negatively associated with senescent (Cdkn2a-positive) and pit-like (Tff1-positive) cells, we analyzed *Ror2* knockout mice for a possible shift in cellular identity. Characterization of Krt19-positive ductal structures at 20 weeks post Tamoxifen revealed that lesions in *Ror2* knockout mice were less proliferative and had a significantly higher number of Cdkn2a-positive senescent cells, with the senescent phenotype having abated by 40 weeks (**Fig. 4B, 4C**). The high prevalence of mucinous lesions at 40 weeks led us to stain for Muc6, a SPEM marker, as well as Tff1 and Muc5ac, both markers for gastric pit cell identity. We observed a slight reduction of *Muc6*-expressing Krt19-positive cells (**Fig. 4D**), accompanied by a strikingly higher proportion of ductal cells expressing Muc5ac and Tff1 when *Ror2* was ablated (**Fig. 4E, 4F**). This shift of cell identity also was evident in immunoblot analyses from bulk pancreatic tissue lysates confirming increased Tff1 levels in *Ror2*-*null*, *Kras^G12D^*-expressing mice (**Fig. 4G**).

Altogether, these data show that ablation of Ror2 counteracts a wound-healing response as indicated by the presence of less proliferative meta/dysplastic cells, while promoting tissue transformation and fibrosis. Ablation also had a profound impact on cellular identity of precancerous lesions, favoring differentiation towards a mucinous Tff1-positive gastric pit cell-like identity, with an overall higher likelihood of progression to PDAC. These two concurrent phenotypes suggest a possible connection between the pit-like phenotype and progression to carcinoma, a possibiltiy bolstered by Tff1 also being a well-established marker for the classical PDAC molecular subtype [23, 25].

### Pit cell identity marks the classical subtype of PDAC whereas Ror2 expression is associated with the basal-like subtype

To assess if Ror2 and gastric gene signatures define subtype identity in PDAC, we turned to the well-established KPC (*Ptf1aCre;Kras^LSL-G12D^;Trp53^R172H^*) mouse model, which recapitulates human PDAC subtype heterogeneity including the epithelial classical and the more mesenchymal, basal-like subtypes [40]. Dual IHC for Ror2 (brown) and Tff1 (teal) revealed that expression is very heterogeneous in neoplastic and carcinoma areas in KPC mice and confirmed that Ror2 and Tff1 define distinct, non-overlapping subpopulations (**Fig. 5A**). Accordingly, we also found a significant negative correlation among the two genes in RNA-seq data previously generated from mouse pancreatic cancer cell lines [41]. Moreover, while *Ror2^Low^/Tff1^High^* cells are characterized by a differentiated epithelial cell identity (clusters C2a-C2c), *Ror2^High^/Tff1^Low^* expressors are associated with a more mesenchymal cell differentiation state (C1) (**Fig. 5B**).

**Fig. 5:**
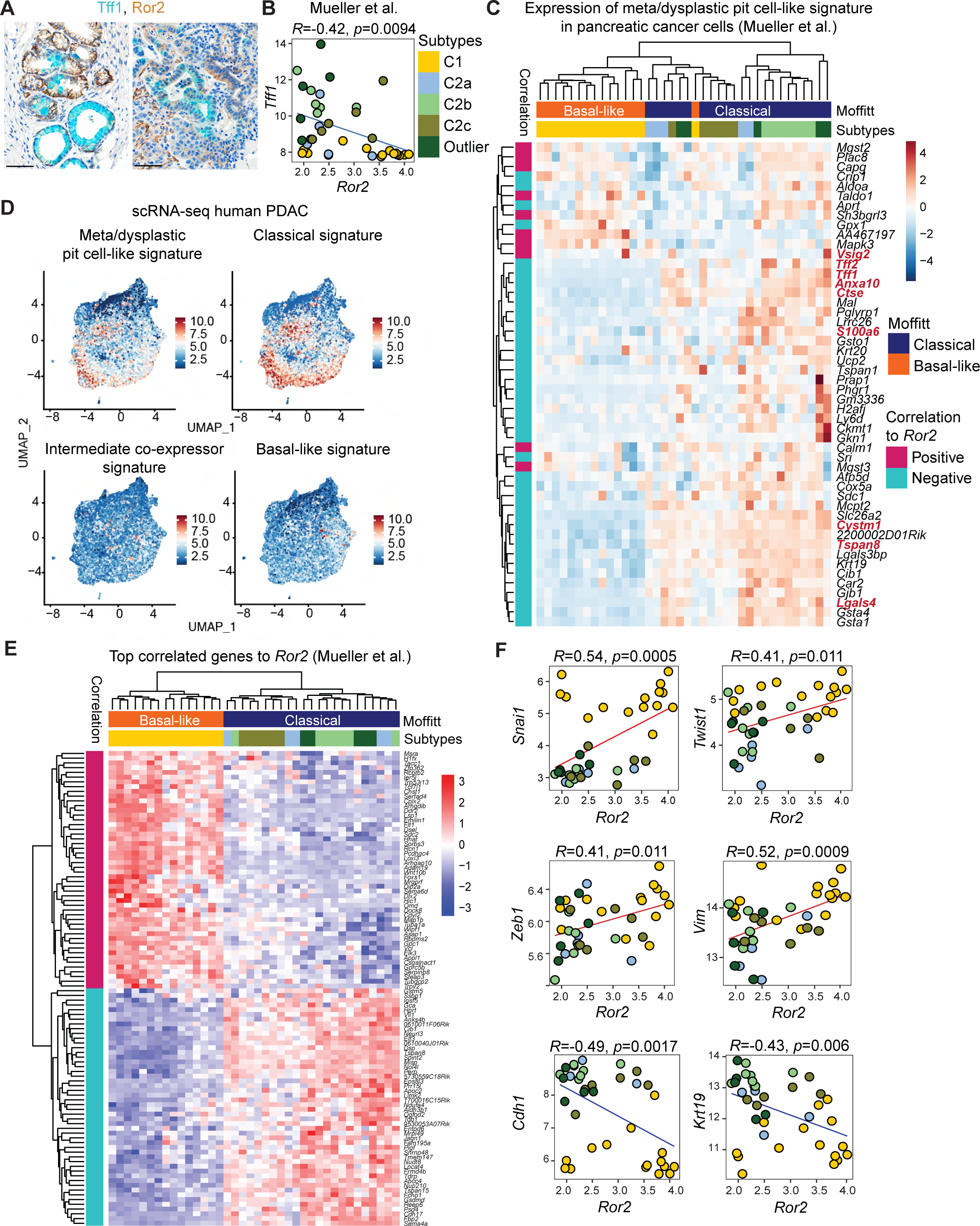
(**A**) Representative images of dual IHC staining for Tff1 (teal) and Ror2 (brown) on tissue of *KPC* mice. Scale bars, 50 µm. (**B**) Pearson correlation scatter plots showing correlation of *Ror2* to *Tff1* gene expression in pancreatic cancer cell lines. RNA-seq data from Mueller et al., correlation coefficient and p-value are indicated. (**C**) Heatmap depicting expression of top 50 most enriched genes in meta/dysplastic pit-like cells (Schlesinger et al.) in pancreatic cancer cell lines (Mueller et al.). PDAC subtype annotation was performed with gene signatures from Moffitt et al.. (**D**) UMAP plots indicating expression signatures of meta/dysplastic pit-like cells and indicated PDAC subtypes retrieved from Ragharan et al. in 174 human scRNA-seq PDAC samples. (**E**) Heatmap using RNA-seq data from Mueller et al., showing expression of top 50 genes that most significantly positively or negatively correlate with *Ror2*. For subtype annotation, gene signatures from Moffitt et al. were used. (**F**) Correlation of *Ror2* expression to indicated genes was determined by Pearson correlation. RNA-seq data from Mueller et al. was used.

Using Pearson correlation analysis, we further uncovered that *Ror2* does not only negatively correlate to *Tff1*, but to a myriad of genes that define meta/dysplastic pit-like cells. Importantly, the great majority of most enriched pit cell-like marker genes (n=50) show a strong association to the classical subtype, including 9 genes (highlighted in red) that are part of a recently defined 30 gene human classical PDAC signature [26] (**Fig. 5C**). To validate these findings in human PDAC tissue, we surveyed publicly available scRNA-seq data from 174 human PDAC samples, to our knowledge, the largest aggregated collection of PDAC patient tissues [2, 42–50]. Ductal cells (KRT18 and KRT19 enriched, negative for acinar and other cell lineage markers) were subsetted from total cells and re-clustered in Seurat. GSEA within the ductal population revealed a tight overlap between the pit cell-like and the classical PDAC gene signature (**Fig. 5D, Suppl. Fig. 5A**). These data suggest that cell fate traits of meta/dysplastic pit-like cells are maintained in classical PDAC in mouse models and importantly, in patient tumors.

To identify cellular identity features that show high correlation to *Ror2*, we re-visited the RNA-seq data of mouse pancreatic cancer cell lines [41] given its large number of samples that represent classical or basal-like PDAC subtype identity. Strikingly, genes that show highest correlation to *Ror2* expression exhibit markedly elevated expression in the mesenchymal cell lines (C1) associated with the basal-like subtype, as defined by gene signatures from Moffitt et al. [22] (**Fig. 5E**). To elucidate the biological function of genes that positively and negatively correlated with *Ror2*, we conducted GO pathway analysis. Positively-correlating genes were associated with cancer cell aggressiveness and EMT, whereas negatively-correlating genes had a role in apoptosis or epithelial differentiation (**Suppl. Fig. 5B**). Interestingly, the most significantly enriched GO term for positively-correlating genes “multicellular organism development” contained a myriad of well-known EMT-inducing transcription factors, such as Prrx1, Snai, Twist or Zeb family members. Correlation plots present significant positive correlation of *Ror2* with *Snai1*, *Twist1*, *Zeb1* or *Vimentin (Vim)* expression, while expression of epithelial genes *E-Cadherin (Cdh1)* and *Krt19* is negatively correlated with *Ror2* (**Fig. 5F**). Interestingly, *Ror2*-positive pancreatic cancer cells were associated with lower *Krt19* expression, a finding that we observed previously in Ror2-positive precancerous lesions.

### ROR2 induces EMT and stemness in human pancreatic cancer cells

Based on our finding that Ror2 correlated to the basal-like/mesenchymal PDAC subtype in mice, we asked if ROR2 showed a similar association in human PDAC. For this, we examined ROR2 expression in resected tumors, patient-derived organoids and cancer cell lines. IHC of human pancreas tissue revealed that ROR2 is expressed at low levels in normal tissue, while a proportion of precancerous lesions and PDAC cells are strongly positive (**Fig. 6A**). Analysis of RNA-seq data from micro-dissected, epithelial proportions of human PDAC samples previously published by Maurer et al. [51] revealed, that *ROR2* expression significantly correlates with the expression of EMT genes, such as *ZEB1* and *Vimentin*, whereas *E-Cadherin* and *EPCAM* inversely correlate (**Fig. 6B**). Of note, basal-like tumor samples are highly underrepresented in human PDAC, likely due to aggressive, invasive tumors not usually qualifying for resection. Accordingly, when we analyzed human organoid (**Suppl. Fig. 6A)** and pancreatic cancer cell lines, including primary patient lines, we found that the majority exhibited more epithelial characteristics and were negative for *ROR2* expression. However, all lines that did express ROR2 also showed mesenchymal features, such as Vimentin and ZEB1 expression, and extremely low levels of the E-Cadherin (**Fig. 6C, 6D**). We conclude that ROR2 is indeed associated with a mesenchymal differentiation in human PDAC.

**Fig. 6:**
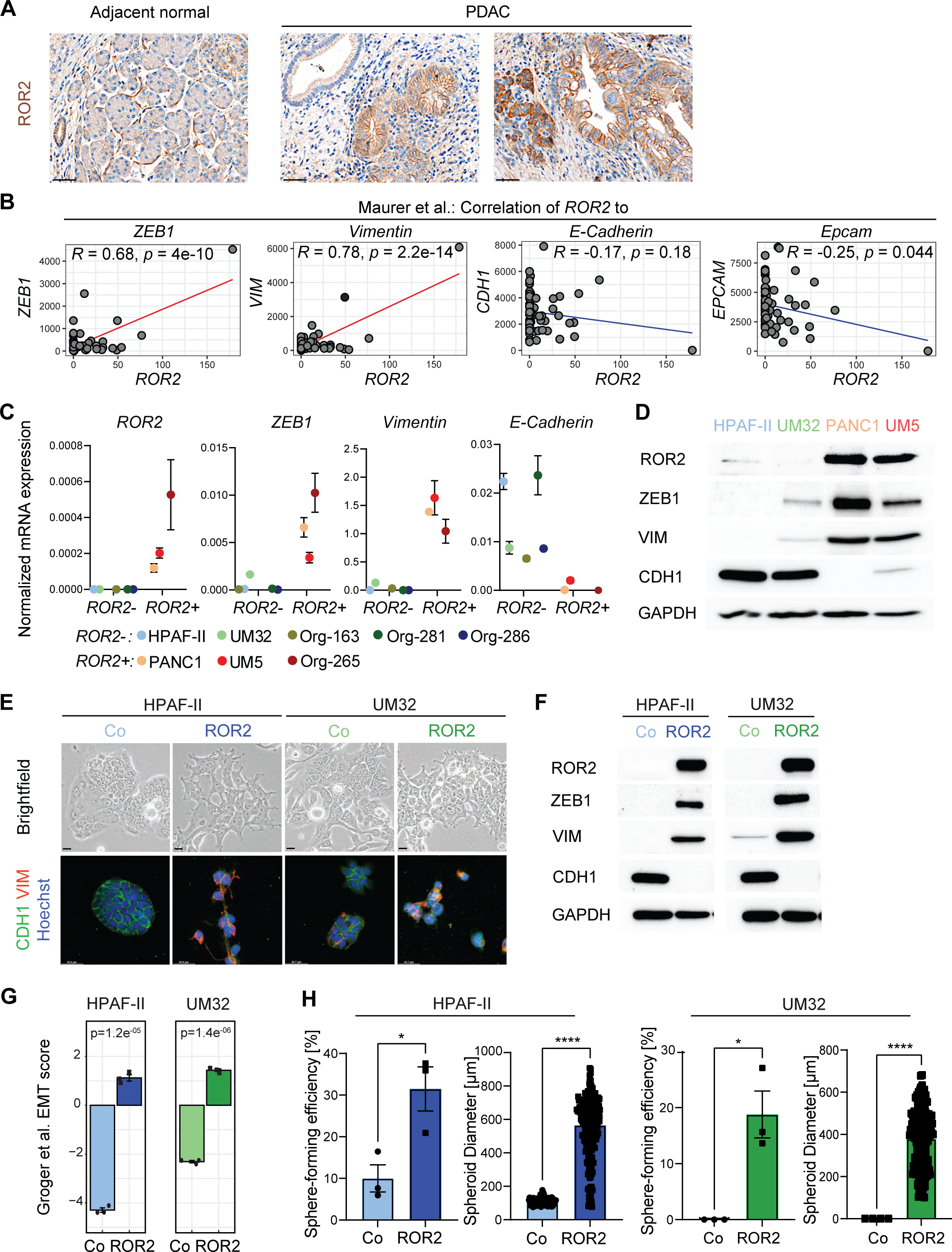
(**A**) Representative brightfield images of ROR2 IHC staining on human PDAC tissue. Scale bars, 50 µm. (**B**) Correlation plots showing correlation of *ROR2* mRNA expression to indicated genes in micro-dissected, epithelial PDAC compartment using RNA-sequencing data from Maurer et al.. (**C**) mRNA expression of *ROR2*, *ZEB1*, *VIM*, and *CDH1* was determined in indicated human PDAC and organoid lines by performing qRT-PCR. Expression values were calculated in relation to the housekeeper gene *GAPDH* (n=3). (**D**) Representative immunoblot analysis of indicated proteins using protein lysates of HPAF-II, UM32, PANC1 and UM5 cells. GAPDH served as loading control. (**E**) Brightfield images of control and ROR2-overexpressing HPAF-II and UM32 cells (upper panel). Scale bars, 200 µm. Immunofluorescence staining for CDH1 (green), VIM (red) and nuclei (Hoechst 33342, blue) of indicated cell lines (lower panel). Scale bars, 37.6 µm. (**F**) Representative immunoblot analysis detecting ROR2, VIM, ZEB1 and CDH1 protein levels in control and ROR2-overexpressing HPAF-II and UM32 cells. GAPDH served as loading control. (**G**) Transcriptomic signatures of control and ROR2-overexpressing HPAF-II and UM32 cells were compared to EMT score, established by Groger et al.. p-values were calculated by Mann-Whitney test. (**H**) Sphere forming efficiency of control and ROR2-overexpressing HPAF-II and UM32 cells is depicted as number of spheres in relation to total number of seeded wells (n=3). Diameter for every sphere was determined. Data are presented as mean ± SEM; p-values were calculated by two-tailed, unpaired Student’s *t*-test; * p < 0.05, **** p < 0.0001.

Next, we assessed if expression of *ROR2* in the epithelial cell lines HPAF-II and the primary patient cell line UM32 is sufficient to cause cellular reprogramming and a switch of cell identity. Notably, expression of ROR2 clearly induced EMT, characterized by loss of the epithelial and gain of the spindle-shaped, mesenchymal morphology (**Fig. 6E**). Immunofluorescence and immunoblot analyses demonstrated concomitant expression of the EMT markers ZEB1 and Vimentin, while E-Cadherin was completely lost (**Fig. 6E, 6F**). Consequently, we compared RNA-seq data of HPAF-II and UM32 control and ROR2-overexpressing cells to the EMT signature established by Groger et al. [52]. ROR2-overexpressing HPAF-II and UM32 cells displayed a significantly enriched EMT score compared to control cells (**Fig. 6G**). In addition, expression of previously identified classical PDAC subtype and pit cell marker genes were completely abrogated upon *ROR2* expression, overall indicating that ROR2 erodes pancreatic cancer cell identity and promotes acquisition of a mesenchymal phenotype (**Suppl. Fig. 6B**).

Previous studies revealed that activation of EMT programs induce stem cell features, with ZEB1 being a key factor for stemness in various cancer entities [4, 53, 54]. Our RNA-sequencing data unveiled that the gastric stem cell genes *LGR5*, *SOX2* or *CXCR4* are greatly upregulated upon expression of *ROR2* in HPAF-II and UM32 cells (**Suppl. Fig. 6C**). To test if this stem cell-like signature predicts function, we examined the ability of ROR2-expressing cells to form tumor spheres in low adhesion plates. Indeed, ROR2-expressing cells formed a significantly greater number and larger tumor spheres *in vitro* than control cells (**Fig. 6H**). Overall, our data identify ROR2 as a substantial driver of human PDAC progression, inducing EMT and stemness.

### ROR2 confers a RAS-independent, AKT-dependent proliferative phenotype

Next, we studied the effects of ROR2 loss in the ROR2-expressing, mesenchymal PANC1 and UM5 cell lines. CRISPR/Cas9-mediated knockout using two different guideRNAs (ROR2 KO #1, ROR2 KO #2) had no consistent effect on EMT marker gene expression (**Suppl. Fig. 7A**), demonstrating that ROR2 signaling is not necessary for maintaining EMT once initiated. However, ablation of ROR2 in PANC1 and UM5 led to a significant decrease in cell proliferation (**Fig. 7A**), an effect that was particularly dramatic in the patient-derived primary cell line UM5, so much so that recovery of RNA/protein or execution of functional assays was extremely challenging (**Suppl. Fig. 7B**). To study immediate ROR2 target genes, we performed siRNA-mediated knockdown of *ROR2* and performed RNA-seq. Top enriched GSEA gene sets in control versus *ROR2* knockdown cells were associated with fundamental regulators of proliferation, such as cell cycle-regulating E2F targets and G2/M transition genes (**Suppl. Fig. 7C**), an effect that has been observed in other tumor types [55, 56]. In contrast, ectopic overexpression of ROR2 in the ROR2-cell lines HPAF-II and UM32 induced an extensive increase of cell proliferation (**Fig. 7B**), correlating with an up-regulation of E2F and G2/M target genes (**Suppl. Fig. 7D**).

**Fig. 7:**
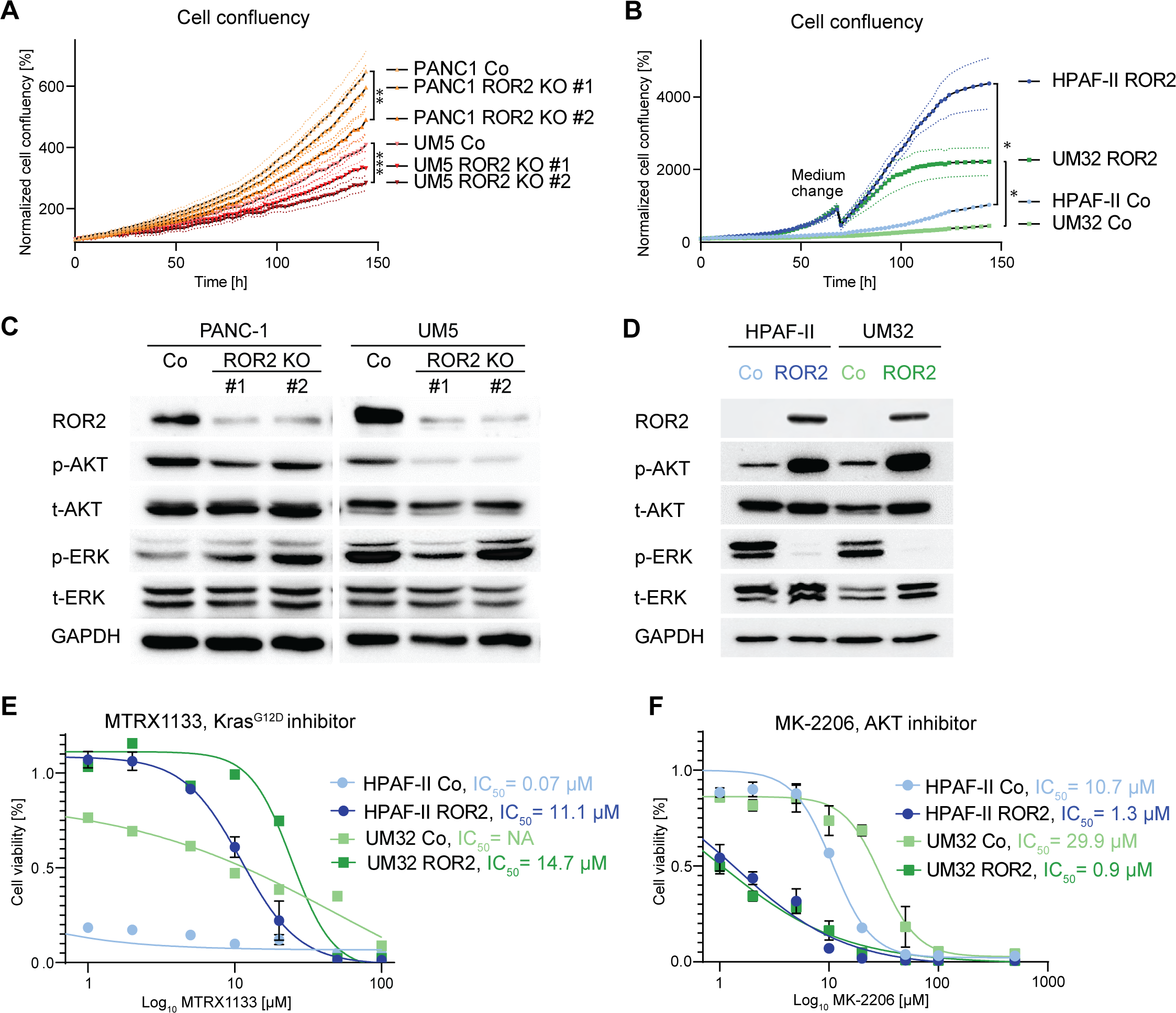
(**A**) Cell confluency of PANC1 and UM5 control and ROR2 knockout cells using two different guide RNAs (ROR2 KO #1, ROR2 KO #2) was measured every 2 hours over a period of 6 days (n=4). Each data point was calculated as relative cell confluency in relation to the first measurement. Statistical significance was determined for the last measurement. (**B**) Cell confluency of control and ROR2-overexpressing HPAF-II and UM32 cells was monitored for a total of 6 days. (**C**) Representative immunoblot analysis for ROR2 and indicated signaling proteins in PANC1 and UM5 control and ROR2 knockout cells. GAPDH served as loading control. (**D**) Representative immunoblot analysis showing expression levels and phosphorylation status of indicated signaling pathway regulators in control and ROR2-overexpressing HPAF-II and UM32 cells. GAPDH served as loading control. (**E**) Control and ROR2-overexpressing HPAF-II and UM32 cells were treated with increasing concentrations of Kras^G12D^ inhibitor MRTX1133 and dose-response curves were established (n=3). IC_50_ was calculated. (**F**) Dose-response curves after treating control and ROR2-overexpressing HPAF-II and UM32 cells with indicated concentrations of AKT inhibitor MK-2206 (n=3). For each cell line, IC_50_ was determined. Unless stated otherwise, all data are presented as mean ± SEM; p-values were calculated by two-tailed, unpaired Student’s *t*-test; * p < 0.05, ** p < 0.01, *** p < 0.001.

We next used exploratory phospho-Arrays to identify downstream signaling of ROR2 (data not shown) and confirmed these findings by immunoblot analysis. While ROR2 knockout cells, especially UM5, showed a reduction in phospho-AKT levels, overexpression of *ROR2* led to a marked increase, accompanied by a virtual ablation of activated phospho-ERK (**Fig. 7C, 7D**), a surprising effect considering both cell lines harbor KRAS^G12D^ mutations. To test if this profound shift away from ERK and towards AKT signaling would alter response to pharmacological inhibitors, we tested HPAF-II and UM32 ROR2 overexpressors’ response to the KRAS^G12D^ inhibitor MRTX1133 (**Fig. 7E**) and the AKT inhibitor MK-2206 (**Fig. 7F)**. Indeed, we found that ROR2 expressing cells were more resistant to KRAS^G12D^ inhibition compared to controls, but newly vulnerable to AKT inhibition.

## DISCUSSION

Pancreatic cancer is an almost universally fatal disease marked by immense tumor cell heterogeneity that contributes to resistance to therapy [1]. This tumor cell heterogeneity, in turn, is driven by the profound plasticity of the exocrine pancreas [57], a quality evident in acinar cell transdifferentiation to metaplastic duct-like cells in chronic pancreatitis, and in their susceptibility to form precancerous neoplastic lesions induced by the expression of oncogenic Kras [19, 20]. Maintenance of a differentiated cellular identity is inherently at odds with the initial steps of pancreatic cancer formation. Genetic variants in several factors that maintain the differentiated cell identity, including *PDX1*, have been associated with increased susceptibility to PDAC in people [18]. In this study, we have applied single-nucleus ATAC-seq to capture early reprogramming of Kras^G12D^-expressing acinar cells, as well as in acinar cells with a compromised identity due to *Pdx1* ablation, which are highly susceptible to rapid transformation. With this approach we discovered that ablation of *Pdx1* alone initiates reprogramming towards a gastric cell identity, an effect also seen upon ablation of *Ptf1a*, another transcription factor critical for the maintenance of acinar cell identity [58]. Gastric cell identity gene expression was intensified by the expression of oncogenic Kras. Among the upregulated genes we identified was the non-canonical Wnt receptor Ror2, which had not previously been associated with gastric identity, but which we found was expressed in the deep antral cells of the stomach, where many markers of gastric metaplasia (SPEM) are also expressed [59].

Our findings reveal that in neoplasia, Ror2 remains associated with and promotes a proliferative neck cell/SPEM-like identity. In the stomach and the pancreas, SPEM is associated with a wound healing function, suggesting that, in neoplasia, SPEM is part of a repair program responding to the tissue damage induced by transformation [60]. Indeed, ablation of Ror2 in mouse neoplasia showed more extensive transformation, with early neoplastic lesions being highly senescent and in late neoplasia became dominated by a gastric pit cell-like cell identity. Interestingly, a small subset (3/11) of *Ptf1a^ERT^;K*,Ror2^f/f^* mice developed carcinoma at a very early 40-week time point, a progression speed not observed in control *Ptf1a^ERT^;K\** animals either in this study or historically. This leads us to speculate that neoplastic pit-like cells are direct precursors to carcinoma, in which the pit cell signature is initially maintained, and that their hyperplasia upon Ror2 ablation increases the probability of PDAC development.

In PDAC, we found that *ROR2* expression maintains its inverse correlation with the TFF1-positive gastric pit cell signature that marks the classical molecular subtype. *ROR2* expression is more broadly associated with the aggressive basal-like molecular subtype and a mesenchymal phenotype. Its ectopic expression in human PDAC cell lines with a classical subtype identity caused extensive reprogramming towards a TFF1-negative, mesenchymal, and highly proliferative phenotype. As has been observed in other tumor types, ROR2 is an active regulator of EMT [30, 31]. Interestingly, inhibition of TFF1 in a PDAC cell line induced expression of EMT transcription factors [61], possibly explaining part of the mechanism by which ROR2 induces this dramatic phenotypic switch. Though it is possible that the Ror2-expressing Neck/SPEM neoplasia is a precursor to basal PDAC, other than *ROR2* we did not observe substantial expression of other SPEM markers in this subtype. It is perhaps more likely that ROR2 actively drives mesenchymal differentiation in a manner that reflects its role in fibroblasts, where it is consistently expressed.

Overall, our data point to a complex function of ROR2, resisting progression in precancer stages and promoting progression in PDAC, the latter of which is consistent with previous findings in melanoma, gastric or breast cancer, where it was shown to regulate cancer cell aggressiveness, including proliferation, stemness and EMT [30–32, 62]. However, it can also play tumor-suppressive roles, such as inhibition of invasion and metastasis, as is seen in prostate and endometrial cancer, suggesting that ROR2’s function needs to be carefully evaluated in each cancer type before considering its role as a potential therapeutic target [63, 64]. Notably, a recent study revealed that suppression of Ror1, which belongs to the same receptor family as Ror2, reduced pancreatic cancer growth and metastasis [65]. Based on these and our findings and given the receptors’ cell surface location, targeting ROR2 and its family members may be beneficial for PDAC patients. Overall, the potential of the ROR family as novel therapeutic targets has only been discovered recently and researchers have started to develop small molecule inhibitors and monoclonal antibodies [66–68]. Importantly, a phase 2 clinical trial testing for the ROR2 antibody-drug conjugate BA3021 in patients with head neck cell cancer, melanoma and non-small cell lung cancer is currently in progress (NCT03504488) [69], which will provide much needed information in regards to drug target safety and validation.

Cellular plasticity is increasingly recognized as a mechanism of therapeutic resistance in cancer [26, 70]. ROR2 has been shown to induce EMT in several tumor types, though its relationship to ERK [71, 72] and AKT signaling pathways [73, 74] appears to be highly context-dependent. Notably, ROR2 activity is associated with resistance to BRAF inhibition and chemoresistance in melanoma, though through the activation of ERK and the inhibition of AKT [75]. We found that ROR2 expression in two KRAS^G12D^-mutated PDAC cell lines has a dramatic effect on the classic effector pathways downstream of KRAS, virtually eliminating ERK activity and drastically upregulating AKT activity. ROR2’s hyperactivation of AKT was consistent with observations in several tumor types [73, 74]. This remarkable shift away from a dependency on ERK signaling presaged an impressive resistance to KRAS inhibition by MRTX1133, while the hyperactivation of AKT created an increased vulnerability to AKT inhibition by MK-2206. These data suggest that ROR2 expression in aggressive, basal-like PDAC could be predictive for therapeutic usage of AKT versus, or in addition to, Kras inhibitors in PDAC patients.

In summary, we found that erosion of pancreatic acinar cell identity induced by Pdx1 ablation was sufficient to confer a gastric identity gene signature that was maintained up to and including progression to the classical subtype of PDAC. Among the neck cell/SPEM subpopulation of metaplastic epithelium, we identified Ror2 to be a powerful regulator of cellular identity throughout progression. In early stage disease, Ror2 suppresses transformation and progression to PDAC. In carcinoma, ROR2 promotes progression to an aggressive basal-like subtype where it strongly impacts therapeutic response.

## METHODS

### Cell culture

The human pancreatic cancer cell lines PANC1 and HPAF-II were purchased from ATCC, UM5 and UM32 were kindly provided by Dr. Costas Lyssiotis (University of Michigan). Identity of cell lines was confirmed by STR analysis (University of Arizona). PANC1 were cultivated in Dulbecco’s Modified Eagle Medium (DMEM) (10-013-CV, Corning), supplemented with 10% fetal bovine serum (FBS) (F2242, Thermo Fisher Scientific) and 0.5% Gentamicin (15-710-072, Fisher Scientific). HPAF-II cells were cultured in Minimum Essential Medium (MEM, 11-095-080, Fisher Scientific) containing 1% sodium pyruvate (11360070, Thermo Fisher Scientific), 1% non-essential amino acids (11140050, Thermo Fisher Scientific), 10% FBS and 0.5% Gentamicin. UM5 and UM32 were cultivated in RPMI 1640 (10-040-CV, Corning) with 10% FBS and 0.5% Gentamicin. 293FT cells were grown in DMEM supplemented with 1% sodium pyruvate, 1% non-essential amino acids, 10% FBS and 1% Penicillin/Streptomycin. Generally, cells were passaged at a confluency of ∼80% by using Trypsin/EDTA (25200114, Thermo Fisher Scientific). All cell lines were regularly tested for mycoplasma using the MycoStrip-Mycoplasma Detection Kit (Rep-mys-50, Invivogen) and maintained at 37°C in a humidified chamber with saturated atmosphere containing 5% CO_2_.

### Organoid culture

Organoid cultures 163, 265, 281 and 286 were previously established from PDX tumors [76]. Organoids were maintained in Cultrex^TM^ UltiMatrix Reduced Growth Factor Basement Membrane Extract (BME001-10, R&D Systems) and incubated in Advanced DMEM/F12 (12-634-010, Thermo Fisher Scientific) culture medium, supplemented with human recombinant fibroblast growth factor-basic (FGF2) (100-18C, Peprotech), B27 (17504044, Thermo Fisher Scientific) and other factors [77]. Culture medium was replaced every 2-3 days, and organoids were passaged every 12-14 days. For passaging, organoids were digested in 1 mg/ml collagenase-dispase solution (LS004106, Cedarlane) for 1.5 hours followed by 15 minutes TrypLE^TM^ (12604021, Thermo Fisher Scientific) treatment. Dissociated organoid cells were washed, resuspended in Cultrex and seeded at a density of 20,000 cells per well of a 24 well-plate. After solidification, 1 mL of culture medium was added to each well.

### Gene editing / Overexpression experiments

ROR2 knockout in PANC1 and UM5 cells was induced by using the eSpCas9-LentiCRISPRv2 vector system [78] containing the following guide RNA sequences: ROR2 guide-RNA (gRNA) 1 (AGCCGCGGCGGATCATCATC), ROR2 gRNA 2 (ATGAAGACCATTACCGCCAC) or scrambled control gRNA (ACGGAGGCTAAGCGTCGCAA). For the generation of lentiviruses, 293FT cells were transfected with the target vector and the packaging plasmids pLP1, pLP2 and pLP/VSVG (Thermo Fisher Scientific) by using Lipofectamine 2000 (11668019, Thermo Fisher Scientific). Lentiviral particles were collected 72 hours after transfection and concentrated by using Lenti-X Concentrator (631231, Takara). After transduction, PANC1 and UM5 cells were constantly kept in cell culture medium containing 5 µg/ml and 2.5 µg/ml Puromycin (P8833, Millipore Sigma), respectively. ROR2 knockout was confirmed by Western Blot.

For overexpression of ROR2, pLV[Exp]-EGFP:T2A:Puro-EF1A>hROR2 (VB900004-2980bun,_VectorBuilder) or control vector pLV[Exp]-EGFP/Puro-EF1A>ORF_stuffer, (VB900021-9574dpn, VectorBuilder) were packaged into lentiviral particles, concentrated supernatant was added to HPAF-II and UM32 cells and cells were selected with Puromycin.

### siRNA transfection

One day before siRNA transfection, cells were seeded in antibiotics-free cell culture medium. Cells were transfected with either 10 nM Silencer™ Select Negative Control #1 (4390843, Thermo Fisher Scientific), siROR2 #1 (s9760, Thermo Fisher Scientific) or siROR2#2 (s532077) resuspended in Opti-MEM I Reduced Serum Medium (31985062, Thermo Fisher Scientific) containing Lipofectamine2000. After six hours of incubation, cells were washed with PBS and cultivated in normal cell culture medium. Cells were harvested three and four days after transfection for RNA and protein isolation, respectively. Knockdown efficiency was ∼70%.

### Proliferation assay

To estimate cell proliferation, 2,000 cells were seeded on a well of a 96-well plate and brightfield images were taken every two hours by using the BioTek BioSpa 8 Automated Incubator and Citation 5 reader (Agilent). BioTek Gen5 imaging software (Agilent) was used to determine cell confluency, which is generally represented as the mean of four wells.

### Sphere formation assay

Spheroid formation was studied according to the clonal assay principle. For each experiment, single cells were seeded in wells of three, U-bottom shaped 96-well ultra-low attachment plates (CLS7007-24EA, Millipore Sigma). Cells were cultivated in serum-free RPMI 1640 medium containing B-27 supplement, 10 ng/ml FG2, 20 ng/ml human recombinant epidermal growth factor (EGF) (AF-100-15, Peprotech), 10 mM HEPES (15-630-106, Thermo Fisher Scientific) and 0.5% Gentamicin. Spheroid cultures were kept for 14 days, number of spheres in relation to total number of seeded wells was assessed and sphere forming efficiency was calculated. Only spheres with a diameter > 75 µm were considered. Brightfield images were taken with the THUNDER Microscope Imager (Leica).

### Drug response assay

For drug response assays, 4,000 cells/well were seeded in 100µl of appropriate media in a 96-well plate. After overnight incubation, cells received treatment with increasing concentrations of the Kras^G12D^ inhibitor MRTX1133 (E1051, Selleckchem) and AKT inhibitor MK-2206 (S1078, Selleckchem). All vehicles were treated with 1% DMSO. Cell viability was assessed by using CellTiter-Glo 2.0 reagent (G9242, Promega). After 48-72 hours, 100 µl CellTiter-Glo 2.0 reagent was added to each well, followed by incubation of the plate on a shaker for 2 minutes using 120 rpm and 10 minutes incubation at room temperature. Luminescence was measured by using the SpectraMax iD3 (Molecular Devices). The average of each duplicate was calculated and normalized to the vehicle control. IC_50_ values were calculated using GraphPad Prism version 9.

### Mouse lines

The strains *Ptf1aCre^ERT^*, *Pdx1^f/f^*, *Kras^LSL-G12D^* and *R26R^LSL-YFP^* were previously described [13]. *Ror2^f/f^* animals were purchased from the Jackson Laboratory (strain #018354) [79] and bred into *Ptf1aCre^ERT/+^;Kras^LSL-G12D^* animals. To induce *Ptf1aCre^ERT^*-dependent recombination, animals were treated with Tamoxifen (T5648, Sigma). Tamoxifen was dissolved in corn oil and administered by oral gavage, 5 mg daily for a total of five days. All animal procedures were approved by the University of Michigan and Henry Ford Health System Institutional Care and Use of Animal Committees.

### Acinar cell isolation

Acinar cells were isolated from pancreas tissue as previously described [13]. In brief, pancreas tissue was washed three times in Hank’s Balanced Salt Solution (HBSS) (SH3026801, Thermo Fisher Scientific), minced and digested in HBSS containing 0.5 mg/ml Collagenase P (Roche, 11249002001) for 15 minutes at 37°C. Digestion reaction was stopped by washing the tissue three times in HBSS with 5% FBS. Acinar cells were isolated by straining the tissue through 500 µm and 100 µm filter meshes and centrifugating the cells through a gradient of 30% FBS in HBSS. Acinar cells designated for RNA-sequencing experiments were washed in PBS, resuspended in RLT buffer and RNA was immediately isolated.

### PCR genotyping of Ror2^NULL^ allele

PCR-based genotyping of the *Ror2^NULL^* allele was performed according to Ho et al. [79]. In brief, acinar cells of *Ptf1a^ERT^;K\** and *Ptf1a^ERT^;K*;Ror2^f/f^* mice were isolated one week after Tamoxifen administration and genomic DNA was isolated using the DNeasy® Blood and Tissue Kit (69504, Qiagen). PCR was performed using the Go Taq^TM^ Master Mix (PRM7123, Promega).

### Protein extraction and immunoblotting

For Western Blot analysis, cells were seeded at defined densities and lysed in RIPA buffer (10 mM Tris-HCl (pH 7.4), 150 mM NaCl, 0.1% SDS, 1% Na-deoxycholate and 1% Triton X-100) supplemented with protease (A32965, Thermo Fisher Scientific) and phosphatase inhibitor (4906837001, Millipore Sigma). Cell extracts were sonicated for 10 seconds and residual debris was pelleted by centrifugation (10 min, 18 400 g, 4°C). For protein extraction from pancreata, ∼ 30 mg tissue was added to tissue lysis buffer (10 mM Tris-HCl (pH 7.4), 150 mM NaCl, 1 mM EDTA, 1% NP-40, 0.1% SDS, 0.5% Na-deoxycholate supplemented with protease and phosphatase inhibitors). Tissue was homogenized by using the Qiagen tissue lyser (3 min, 50 oscillations/s). Samples were sonicated and supernatant was extracted after centrifugation (20 min, 18 400 g, 4 °C).

Protein concentration was quantified by using the Pierce BCA Protein Assay Kit (23227, Thermo Fisher Scientific). Protein lysates were denatured at 95 °C for 5 min and 20 - 30 µg of protein were generally loaded on polyacrylamide gels and electrophoresis was performed. Separated proteins were transferred onto a PVDF membrane (IPVH00010, Millipore Sigma) by wet-protein transfer. Membranes were blocked in Tris-buffered saline, 0.1% Tween 20 (TBST) containing 5% milk and primary antibodies as listed in Supplementary Table 1 were applied overnight at 4 °C. The secondary, anti-rabbit HRP-conjugated antibody (45-000-682, Thermo Fisher Scientific) was incubated for one hour at RT and bands were visualized using Clarity or Clarity Max Western ECL substrate (1705061 or 1705062, Bio-Rad) with the ChemiDoc^TM^ Imaging System (Bio-Rad).

### Immunohistochemistry (IHC)

Tissue designated for IHC analysis was fixed overnight in Z-Fix (NC9050753, Thermo Fisher Scientific), washed with PBS for 30 minutes at RT, processed and embedded in paraffin. The use of human samples was approved by the Henry Ford Health Institutional Review Board. IHC single- and multiplex-staining were performed on 4 µm sections using the Discovery ULTRA stainer (Roche). Antibodies and dilutions are provided in Supplementary Table 1. Hematoxylin and Eosin (H&E) and Picrosirius Red staining were performed according to standard protocols. Stained sections were scanned with the PANNORAMIC SCAN II (3DHistech Ltd.) and the number of positively stained cells / area was quantified with HALO software v3.5.3577.214 (Indica Labs). Representative images were taken from scanned images or with the Stellaris 5 confocal microscope (Leica).

### Immunofluorescence (IF)

For IF analysis of adherent pancreatic cancer cells, cells were seeded on chamber slides. Cells were fixed with 4% formaldehyde for 10 minutes at RT, washed with PBS and permeabilized using PBS containing 0.25% Triton X-100 (T8787, Millipore Sigma). After blocking in PBS supplemented with 1% bovine serum albumin (BSA) (BP9703, Thermo Fisher Scientific), 5% donkey serum (D9663, Millipore Sigma) and 0.02% Triton X-100, primary antibodies were incubated overnight at 4°C. Antibodies and dilutions are provided in Supplementary Table 1. Cells were then stained with the secondary antibodies, anti-rabbit 488 (A21206, Thermo Fisher Scientific) and anti-mouse 568 (A10037, Thermo Fisher Scientific) as well as Hoechst 33342 (62249, Thermo Fisher Scientific).

## RNA-ISH

Multiplexed RNA-ISH and IF staining were performed as previously described [80]. In brief, FFPE sections were baked in a dry air oven at 65°C for one hour, deparaffinized and hydrated in xylene and 100% ethanol. After antigen retrieval in a steamer for 15 minutes using the Co-Detection Target Retrieval (323165, Advanced Cell Diagnostics) solution and blocking, the primary antibody was applied. Slides were post-fixed in 10% Neutral Buffered Formalin (NBF) for 30 minutes at RT and treated with RNAscope®Protease Plus (322381, Advanced Cell Diagnostics) at 40 °C for 11 minutes. For application of RNA-ISH probes, the RNA-ISH Multiplex Fluorescent Assay v2 Kit (323110, Advanced Cell Diagnostics) was used according to manufacturer’s instructions. Following RNAscope staining, sections were stained with DAPI at RT for 15 minutes, washed in PBS, secondary antibody was applied and slides were embedded with ProLong Diamond (P36961, Thermo Fisher Scientific) mounting medium.

### Single-nucleus ATAC-seq

Pancreatic tissue was snap-frozen in liquid nitrogen and stored at -80 °C. For isolation of nuclei, frozen tissue was pulverized by using the CP02 pulverizer (Covaris). Pulver was washed with PBS containing 0.04% BSA and 1X protease inhibitor and centrifuged (5 min, 500 g, 4 °C). Supernatant was discarded, pellet was resuspended in 0.1 U/µl DNase solution (EN0521, Thermo Fisher Scientific) and incubated on ice for 5 minutes. Tissue was washed twice with PBS containing 0.04% BSA and 1X protease inhibitor. After the last centrifugation step (5 min, 500 g, 4 °C), supernatant was discarded, pellet was resuspended in freshly prepared lysis buffer (10 mM Tris-HCl, 10 mM NaCl, 3 mM MgCl_2_, 0.1% Tween-20, 0.1% Nonident P40, 0.01% Digitonin (BN2006, Thermo Fisher Scientific), 1% BSA and 1X protease inhibitor) and incubated on ice for 10 minutes. Then, nuclei were isolated by using a 2 ml Dounce homogenizer, 25 strokes with pestle A and 25 strokes with pestle B. To remove connective tissue and residual debris, sample was filtrated through a 70 µm cell strainer (352350, Thermo Fisher Scientific), followed by centrifugation (1 min, 100 g, 4 °C). Supernatant was transferred to a new tube and washed with wash buffer (10 mM Tris-HCl, 10 mM NaCl, 3 mM MgCl_2_, 0.1% Tween-20, 1% BSA and 1X protease inhibitor) followed by centrifugation (5 min, 500 g, 4°C) for three times. Before the final washing step, suspension was filtered through a 40 µm Flowmi cell strainer (BAH136800040-50EA, Millipore Sigma). Finally, nuclei were resuspended in Nuclei Buffer (10X Genomics). Library preparation and sequencing was performed at the Advanced Genomics Core of the University of Michigan. First, single nucleus suspensions were counted on the LUNA Fx7 Automated Cell Counter (Logos Biosystems). Then, single nucleus ATAC libraries were generated using the 10x Genomics Chromium instrument following the manufacturer’s protocol. Final library quality was assessed using the LabChip GX (PerkinElmer). Pooled libraries were subjected to paired-end sequencing according to the manufacturer’s protocol (Illumina NovaSeq 6000). 10,000 cells were profiled with a sequencing depth of 50,000 reads per cell. Bcl2fastq2 Conversion Software (Illumina) was used to generate demultiplexed fastq files.

### Single-nucleus ATAC-seq data processing

Doublet removal was performed using scDblFinder (excluding cells with a -log_10_(q-value)) [81] and AMULET (doublet score > 0.975) [82]. In addition, nuclei were excluded if number of reads with low mapping quality was > 40,000, percent of mitochondrial reads was > 10%, percentage of reads in peaks was < 20%, nucleosome signal was > 0.63, activity was detectable in < 3,200 features or > 32,000 features and TSS enrichment score < 2.5. Next, features with < 500 counts across all samples or with reads in < 1% of cells were excluded. Gene activity scores and features were created using the GeneActivity function from the Signac R package [28]. Genes with an activity score of < 200 across all cells (per-sample) or with activity scores in < 1% of cells (per-sample) as well as nuclei with activity scores in < 1,000 genes were removed. For each nucleus, gene activity scores were normalized and natural log transformed. Uniform Manifold Approximation and Projection (UMAP) and shared nearest-neighbor graph (SNN) were constructed using the Seurat R packages RunUMAP and FindNeighbors functions, respectively [83]. Dimensional reduction was performed via latent semantic indexing (lsi) using dimensions 2:30. Cell clusters were identified using the FindClusters function with smart local moving (SLM) algorithm (resolution=0.4) [84]. UMAP projection created a central cluster of low-quality nuclei that demonstrated comparatively poorer QC metrics, such as significantly lower percentage of reads in peaks, lower TSS enrichment and mapping quality scores. This cluster as well as clusters with < 20 cells were removed. Due to a lymph node contamination in one of the two-week snATAC-seq samples, sequencing of one sample was repeated. The minor batch effect was remedied via ComBat [85] using adjusted cluster labels in the model matrix. UMAP projects were recreated with batch-corrected data as described above. The Seurat R package FindAllMarkers function was used to identify differentially accessible peaks among cell clusters [83]. Potential marker genes were required to be present in at least 25% of cells per cluster and show on average, at least a 0.25-fold difference (log-scale) among cell populations. Based on identified cluster-specific marker genes, clusters were manually annotated.

### RNA isolation and quantitative real-time PCR (qRT-PCR)

For total RNA extraction, the RNeasy® Plus Mini Kit (74134, Qiagen) was used following manufacturer’s instructions. Genomic DNA was digested on column by using the RNase-Free DNase Set (79254, Qiagen). mRNA was reverse transcribed to cDNA with the iScript(tm) cDNA Synthesis Kit (1708891BUN, Bio-Rad). qRT-PCR was performed on the QuantStudio 6 Pro qRT-PCR cycler (Thermo Fisher Scientific) using 20 ng cDNA, 500 nM forward and reverse primers as well as 1X Fast SYBR Green Master Mix per reaction (4385612, Thermo Fisher Scientific). Primer sequences are provided in Supplementary Table 2. Relative gene expression was determined in relation to expression of the housekeeping genes *GAPDH* and *Hprt1* and calculated according to the ΔΔCt method [86].

### Bulk RNA-seq

Total RNA was at least analyzed in triplicates. Library preparation and sequencing of RNA samples from isolated mouse acinar cells were performed at the University of Michigan Advanced Genomics Core. Libraries underwent paired-end sequencing with 37.5M reads per sample. Human RNA samples were processed at MedGenome Inc. RNA-seq library preparation was performed by using the Illumina TruSeq stranded mRNA kit. RNA-seq libraries were paired-end sequenced (PE100/150) with 40M reads per sample.

### Bulk RNA-seq data processing

Sequencing reads of mouse RNA-seq experiments from isolated acinar cells were assessed for quality and quantity metrics using FastQC and multiQC [87] before and after trimming with Trimmomatic [88]. Reads were aligned to the mouse reference genome (version GRCm39.108) using Salmon [89]. Sample-specific transcripts per kilobase million (TPM) estimates were manually converted to gene-specific TPM estimates by summing transcript TPMs per gene. Genes and their respective transcripts were collected using the tr2g_ensembl function from the BUSpaRse R package. The original sample by gene count matrix (TPM) was first adjusted for per-sample sequencing library inspired by DESeq2 [90]. The geometric mean was calculated for each gene across all samples. Gene counts were divided by this geometric mean, resulting in gene count ratios. A sample-specific-median or “sample size factor” of these ratios was finally divided from each gene count per sample. Features with estimates < 1 TPM were removed, gene counts were log_2_ transformed and the top 2,000 most variable genes were selected.

For RNA-seq experiments of human pancreatic cancer cell lines, adapter trimming and removal of ribosomal and mitochondrial genome was carried out with fastq-mcf, cutadapt and Bowtie2 [91], respectively. Reads were mapped to the hg19 genome using STAR [92] and raw read counts were estimated with HTSeq (v0.11.2) [93] and normalized with DESeq2 [90].

### Pseudo bulk ATAC- and RNA-seq (Bi-Omic) cross-correlation analysis

In order to compare the snATAC-seq data to bulk RNA-seq data of isolated acinar cells, a sample by gene *pseudo*-*bulk* count matrix was created. First, UMAP clusters identified as acinar cells were subset from the snATAC-seq Seurat object. Then, acinar cells were grouped by sample and the trimmed arithmetic mean of raw counts was calculated to represent the *pseudo bulk* counts; a fraction of 0.05 of observations was trimmed from each end of per-gene counts before the mean was computed. The *pseudo*-*bulk* sample by gene read count matrix (8×20227) was adjusted for per-sample sequencing library inspired by DESeq2 [90] and calculated as described above. Genes with < 1 count across all samples were removed, gene counts were log_2_ transformed and the top 2000 most variable genes were selected. The *pseudo-bulk* ATAC-seq sample-by-gene (8×2000) transformed count matrix was compared to the RNA-Seq sample-by-gene (12×2000) transformed count matrix. Both matrices were adjusted for per-sample sequencing library and log_2_ transformed. Any missing values were imputed with a value of zero. After performing a sample-by-sample spearman correlation and a principal component analysis on the 374 shared genes across the 12 samples of the RNA-seq matrix (biological replicates per genotype n=3), the most distant samples per experimental condition were removed. Both matrices including a total of 8 samples were then standardized by centering and scaling for each gene using (E-mean(E))/sd(E). ATAC-seq genes were measured for their correlation (Spearman) to their respective RNA-seq genes. Genes that demonstrated a correlation ≥ 0.5 were subset (n=117).

### Analysis of publicly available transcriptome datasets

Single-cell gene expression data from mouse pancreas cells were downloaded from Gene Expression Omnibus (GEO) (GSE141017, Samples GSM4293545-GSM4293554) [19]. Scanpy version 1.9.3 [94] was used for re-processing of the digital count matrixes. Counts were normalized to a target sum of 10,000 after masking highly expressed genes, log-transformed, scaled, and the first 50 principal components were used for Leiden clustering and UMAP projection. The PCA space was batch corrected using Harmony [95] with the sample batch as a covariate and the diversity clustering penalty parameter (theta) = 4, and otherwise default settings. Coarse clustering was achieved with a low Leiden resolution score (0.1), and epithelial cell types were selected after screening clusters for expression of the following marker genes: epithelial (*Krt8*, *Krt18*, *Krt19*), endothelial (*Cdh5*, *Pecam1*, *Plvap*), mesenchymal (*Col3a1*, *Dcn*, *Sparc*) and immune (*Tyrobp*, *Cd52*, *Ptprc*). Samples were then re-processed with the same settings as above. Epithelial cells were again coarsely clustered, and the primary acinar cell cluster was distinguished from ductal-type cells by the presence of exocrine marker genes (*Ctrb1, Prss2, Try5*). Meta/dysplastic cells were identified by tdTomato expression. In brief, K-means (K=2, scikit-learn v1.2.2) clustering was carried out on the log-normalized tdTomato expression levels across all epithelial cells. Cells in the higher expression group that were not present in the predominant acinar cell cluster were assigned as meta/dysplastic cells. To calculate cell type scores, gene lists were compiled from Busslinger et al. (Suppl. Table S8) [33] and scored using the Scanpy function scanpy.tl.score_genes().

Single-cell gene expression data from mouse stomach cells were downloaded from GEO (GSE141017, Sample GSM4293556) [19]. Samples were processed as described above, except that coarse clustering was achieved with a resolution score of 0.04 and cells with < 3,000 UMIs were removed.

For the reanalysis of injury-induced ADM dataset (Ma et al.: GSE172380) [20], all YFP+ cells (GSE172380_Cluster+CelltypeLabel_YFP+_all-samples_QCed.csv.gz) were processed with Scanpy using the same pipeline, except cells with < 1,000 UMIs or > 20% mitochondrial content were discarded.

For analyzing RNA-seq data from mouse PDAC cell lines, data (GSE107458) [41] were downloaded from GEO and aligned to mm10. Quality control and normalization was performed and normalized matrix was subjected to Pearson’s correlation to assess correlation of *Ror2* to the expression of other genes. Top 50 positively and negatively correlated genes were selected for further analysis and a heatmap depicting mean normalized expression values was generated using pheatmap. Distribution of normalized expression for candidate markers genes with respect to *Ror2* were plotted. Moreover, expression of top 50 most significantly enriched genes in pit cell-like ADM from Schlesinger et al. data was visualized in a heatmap.

To analyze previously published RNA-seq data of human PDAC samples (GSE93326) [51], processed data were retrieved from GEO and counts were normalized using the normTransform function from DESeq2. Only micro-dissected tumor samples were selected for further analysis. For PDAC tumor subtype annotation, Moffitt’s top-25 gene signature was employed and clustering was performed using ConsensusClusterPlus. Correlation analysis was performed as described above. Analyses were performed using R (version 4.3.1).

For the analysis of human PDAC scRNA-seq data, samples from 174 primary tumors were included (EGAS00001002543, GSE154778, GSE155698, GSE156405, GSE194247, GSE202051, GSE205013, GSE211644, phs001840.v1.p1, PRJCA001063) [2, 42–50]. Post quality control and data harmonization was performed using Harmony [95] and ductal cells based on expression levels of *KRT18* and *KRT19* were selected for further analysis. Using single sample gene set enrichment (ssGSEA) [96], each cell was scored for the expression of previously defined classical, basal-like, intermediate [26] and the established pit cell-like ADM signature (50 genes). Cell specific scores were converted to z-scores for comparison between signatures and cell clusters were classified according to their signatures. Using Pearson’s correlation, the correlations between the classical and pit cell-like ADM signature as well as the basal-like cell and pit cell-like ADM signature were assessed.

### Functional annotation of bulk RNA-seq data

GSEA (Gene Set Enrichment Analysis) was performed using the GSEA software 4.3.2 [97]. For analysis of mouse RNA-seq data, normalized read counts that were centered and scaled by feature were used as input and enrichment analysis was run for generated SPEM signature gene set. GSEA for human cell line RNA-seq data was performed with normalized read counts and data were aligned to the H: hallmark gene sets. Pathway analysis were performed using the Database for Annotation, Visualization and Integrated Discovery (DAVID) [98, 99]. For the generation of GO terms, genes that exhibited a Pearson correlation coefficient > 0.4 and < -0.4, and a p-value <0.01 were used. EMT score was calculated based on log_2_ normalized read counts by using a custom-built function (https://github.com/umahajanatlmu/useful_commands/blob/main/EMT_signature_score.R) incorporating gene signatures from a meta-analysis of 18 independent gene expression studies [52].

### Quantification and statistical analysis

Statistical significance was calculated by two-tailed, unpaired Student’s *t*-test and Mann-Whitney U test as indicated; * p < 0.05, ** p < 0.01, *** p < 0.001, **** p < 0.0001. Statistical analysis was performed with GraphPad Prism 9 software (GraphPad Software Inc.).

### Data availability

Sequencing data will be made available in the GEO archive.

## Acknowledgments

The authors would like to thank the University of Michigan Epigenomics Core and Advanced Genomics Core for tissue preparation and sequencing services, respectively. We would like to thank Linda C. Samuelson and Stephen Parker for the helpful discussions. This work was supported by R01CA235141, U01CA224145 and U01CA274154 (to HCC). SB was supported by a postdoctoral fellowship from the German Research Foundation (DFG, BE 7224/1-1) and a research grant provided by the Sky Foundation.

