## Supplemental Figures for "ROR2 regulates cellular plasticity in pancreatic neoplasia and adenocarcinoma"

#### SUPPLEMENTAL DATA

##### Supplemental figure legends

**Suppl. Fig. 1:** (**A**) Graphical scheme depicting the snATAC-seq workflow. In brief, snap-frozen pancreatic tissue from mice was pulverized and nuclei were isolated by using a Dounce Homogenizer. Tn5-catalyzed DNA tagmentation was performed and followed by microfluidics-based library preparation (10X Genomics), sequencing and data analysis. Graph was created with BioRender.com. (**B**) UMAP depicting low-quality, underperforming nuclei (center location) identified by a low fraction of reads in peaks, suggesting a low signal-to-noise ratio and specificity. (**C**) Heatmap indicating lack of intra-cluster correlation within the low-quality cluster. Inconsistent chromatin accessibility patterns were likely due to underperformance of the tagmentation reaction. (**D**) Heatmap showing expression levels of top 5 most enriched genes in each annotated cell cluster. (**E**) UMAP analyses from all snATAC-seq samples combined. Predicted gene expression of selected marker genes was used for cluster annotation.

**Suppl. Fig. 2**: (**A**) UMAP plots showing predicted gene expression based on gene accessibility of indicated genes in subsetted acinar cell clusters (upper panel). mRNA expression of selected genes in isolated acinar cells from indicated genotypes was determined by qRT-PCR (n=3-4). All data are presented as mean ± SEM; p-values were calculated by two-tailed, unpaired Student's *t*-test; * $p < 0.05$, ** $p < 0.01$, *** $p < 0.001$ and **** $p < 0.0001$. (**B**) UMAP plots for mouse stomach cells (Schlesinger et al.) with gastric cell type annotation guided by Busslinger et al.. (**C**) Dual IHC depicting Ror2 (brown) and Aqp5 (teal) staining in mouse stomach antrum and corpus regions. Scale bars, 50 µm.

**Suppl. Fig. 3:** (**A**) UMAP analyses of meta/dysplastic cells from the Schlesinger et al. scRNA-seq dataset showing enrichment of pit cell signature and *Tff1* expression. (**B**) UMAP plot depicting *Mki67* expression. (**C**) UMAP plot showing *Cdkn2a* expression. (**D**) UMAP plot for *Krt19* expression. (**E**) UMAP plots illustrating cell types found in experimental pancreatitis tissue as well as *Ror2* expression in scRNA-seq dataset from

Ma et al.. (**F**) UMAP plot showing enrichment of SPEM signature. (**G**) UMAP plot showing enrichment of neck cell signature from Busslinger et al.. (**I**) UMAP analyses depicting *Muc6* and *Aqp5* expression.

**Suppl. Fig. 4:** (**A**) PCR for Exon 3 and 4 of the *Ror2* gene, confirming *Cre*$^{ERT}$-mediated targeting of the *Ror2* allele in isolated acinar cells of *Ptf1a*$^{ERT}$*;K*;Ror2*$^{f/f}$ mice compared to those of *Ptf1a*$^{ERT}$*;K** animals. Acinar cells were harvested 1 week after Tamoxifen administration. (**B**) To visualize recombination efficiency, IHC for the reporter YFP as well as Ror2 in *Ptf1a*$^{ERT}$*;K** and *Ptf1a*$^{ERT}$*;K*;Ror2*$^{f/f}$ mice, 20 weeks post Tamoxifen, was performed. (**C**) Representative H&E staining of *Ptf1a*$^{ERT}$*;K** and *Ptf1a*$^{ERT}$*;K*;Ror2*$^{f/f}$ mice, 10 weeks after Tamoxifen. Tissue remodeling was quantified (n=8-9). (**D**) Picrosirius Red staining for mice sacrifized 20 and 40 weeks after Tamoxifen. For all staining, representative images are depicted. Scale bars, 50 µm. Staining were quantified with the HALO software and all data are presented as mean ± SEM; p-values were calculated by two-tailed, unpaired Student's *t*-test; * p < 0.05, *** p < 0.001.

**Suppl. Fig. 5:** (**A**) Pearson correlation plots illustrating correlation of meta/dysplastic pit cell-like signature to ductal cell clusters that showed highest expression of classical and basal-like markers. (**B**) Bar graph depicting selected, significantly enriched GO terms (biological process) from genes that showed a significant correlation or anti-correlation to *Ror2* in pancreatic cancer cell lines (Mueller et al.).

**Suppl. Fig. 6:** (**A**) Representative brightfield images of organoid lines. Scale bars, 100 µm. (**B**) qRT-PCR was used to determine mRNA expression of indicated pit cell marker genes also associated with classical PDAC subtype identity in control and ROR2-overexpressing HPAF-II and UM32 cells. Data are presented as mean ± SEM; p-values were calculated by two-tailed, unpaired Student's *t*-test; ** p < 0.01, ***p < 0.001, **** p < 0.0001. **(C)** Heatmap depicting expression levels of selected EMT and CSC marker genes in control and ROR2-overexpressing HPAF-II and UM32 cells.

**Suppl. Fig. 7:** (**A**) qRT-PCR data showing mRNA expression of *CDH1* and *VIM* in indicated cell lines. Expression values were calculated in relation to the housekeeper gene *GAPDH* (n=3-4). (**B**) Brightfield images of UM5 control and ROR2 knockout cells (ROR2 KO #2) from start to six days of cultivation. (**C**) GSEA analysis showing enrichment of genes regulating E2F-dependent cell cycle regulation and G2/M transition in UM5 control versus siROR2-treated cells. (**D**) GSEA plot depicting enrichment of genes regulating E2F-dependent cell cycle regulation and G2/M transition in UM32 ROR2 overexpressing versus control cells. Unless stated otherwise, all data are presented as mean ± SEM; p-values were calculated by two-tailed, unpaired Student's *t*-test; * $p < 0.05$, ** $p < 0.01$.

#### Supplementary Figure 1

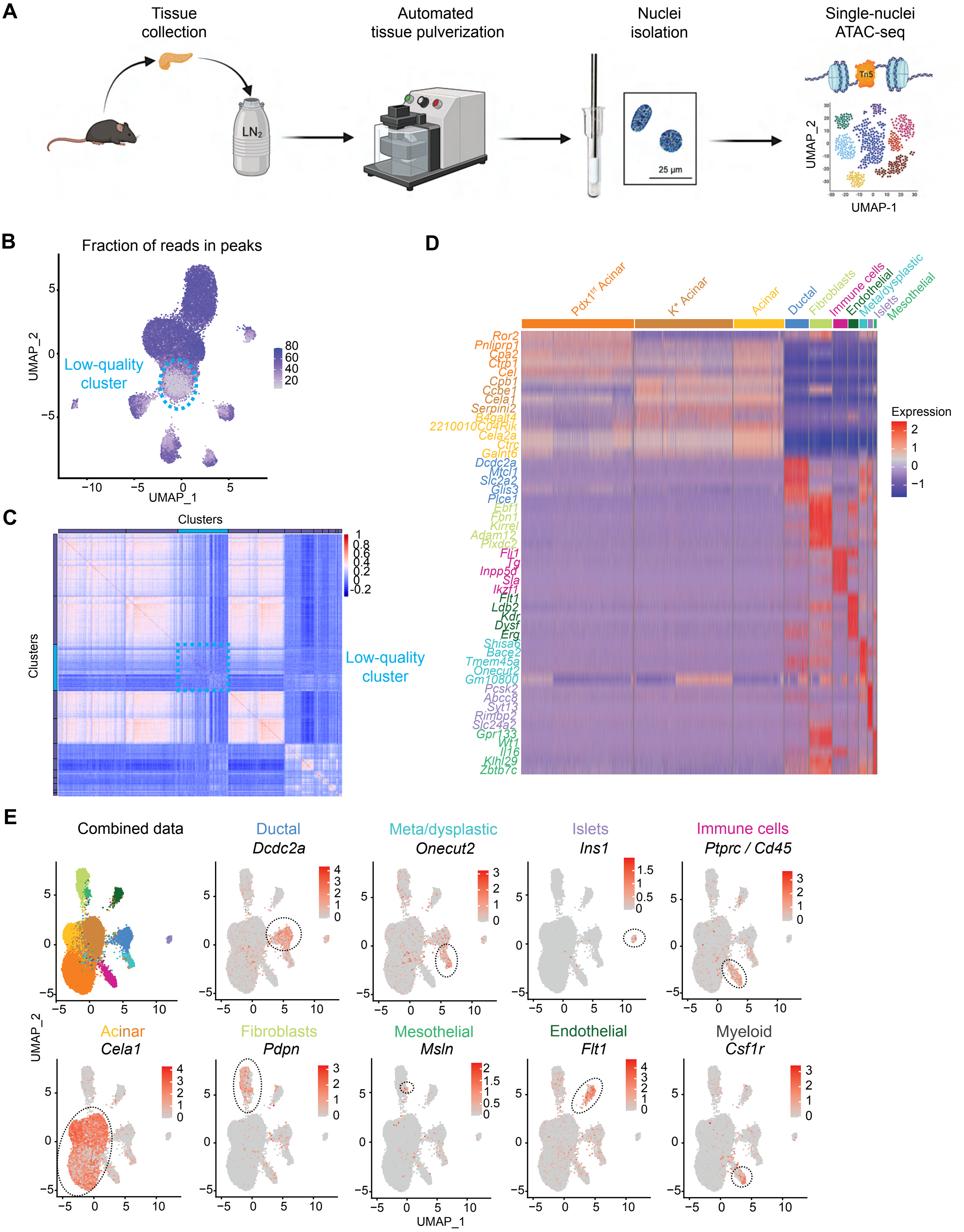

### Supplementary Figure 2

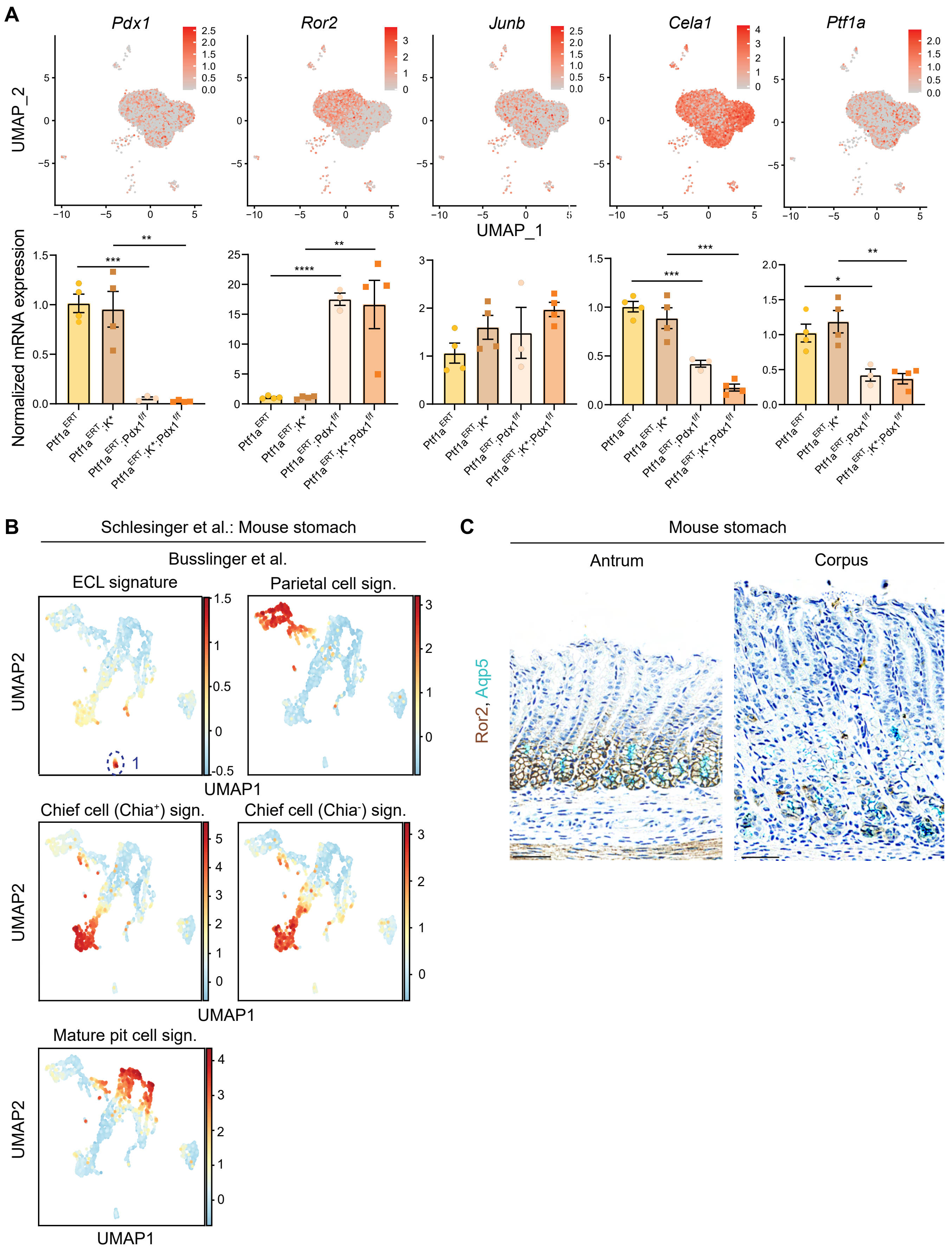

#### Supplementary Figure 3

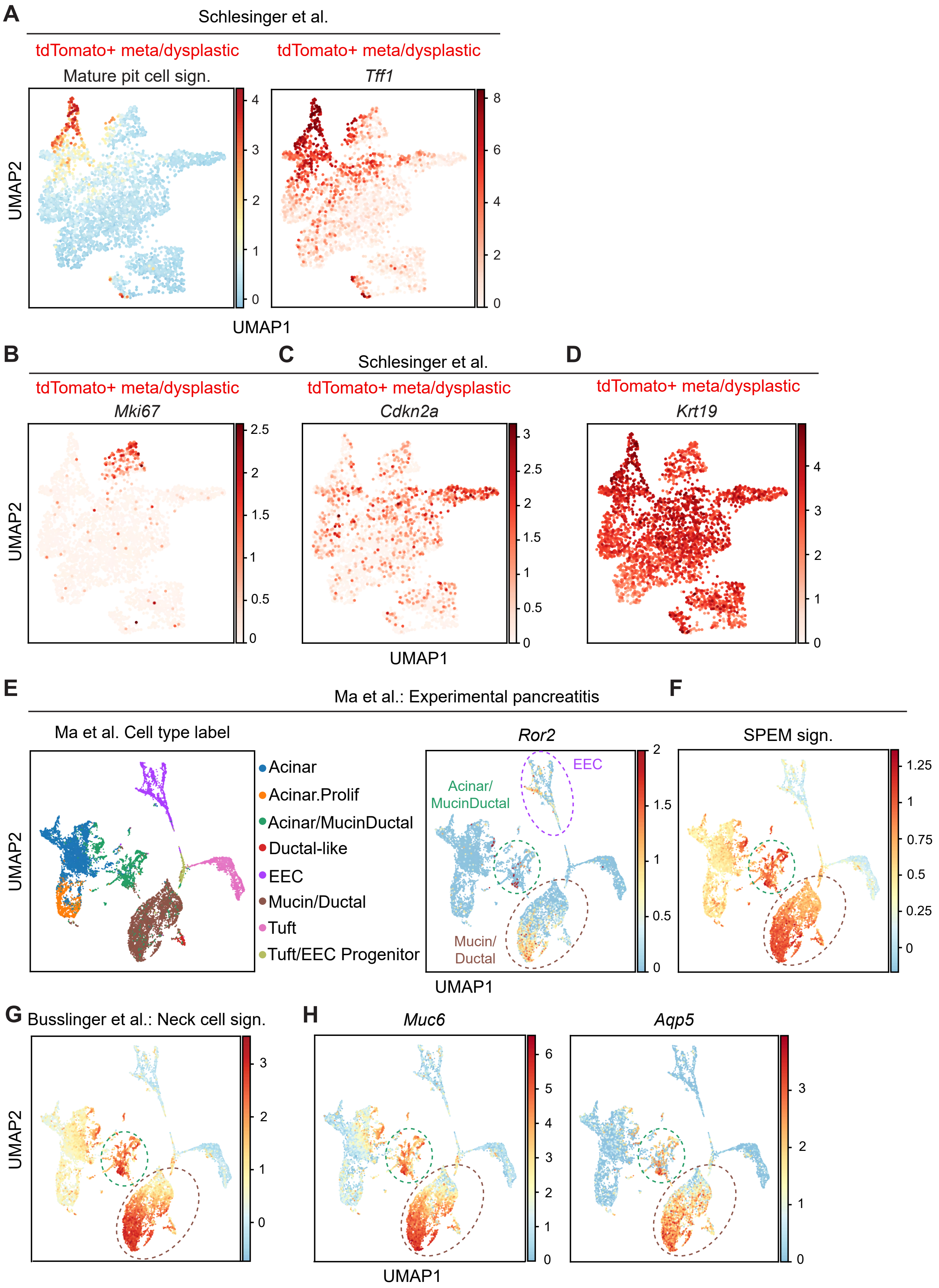

#### Supplementary Figure 4

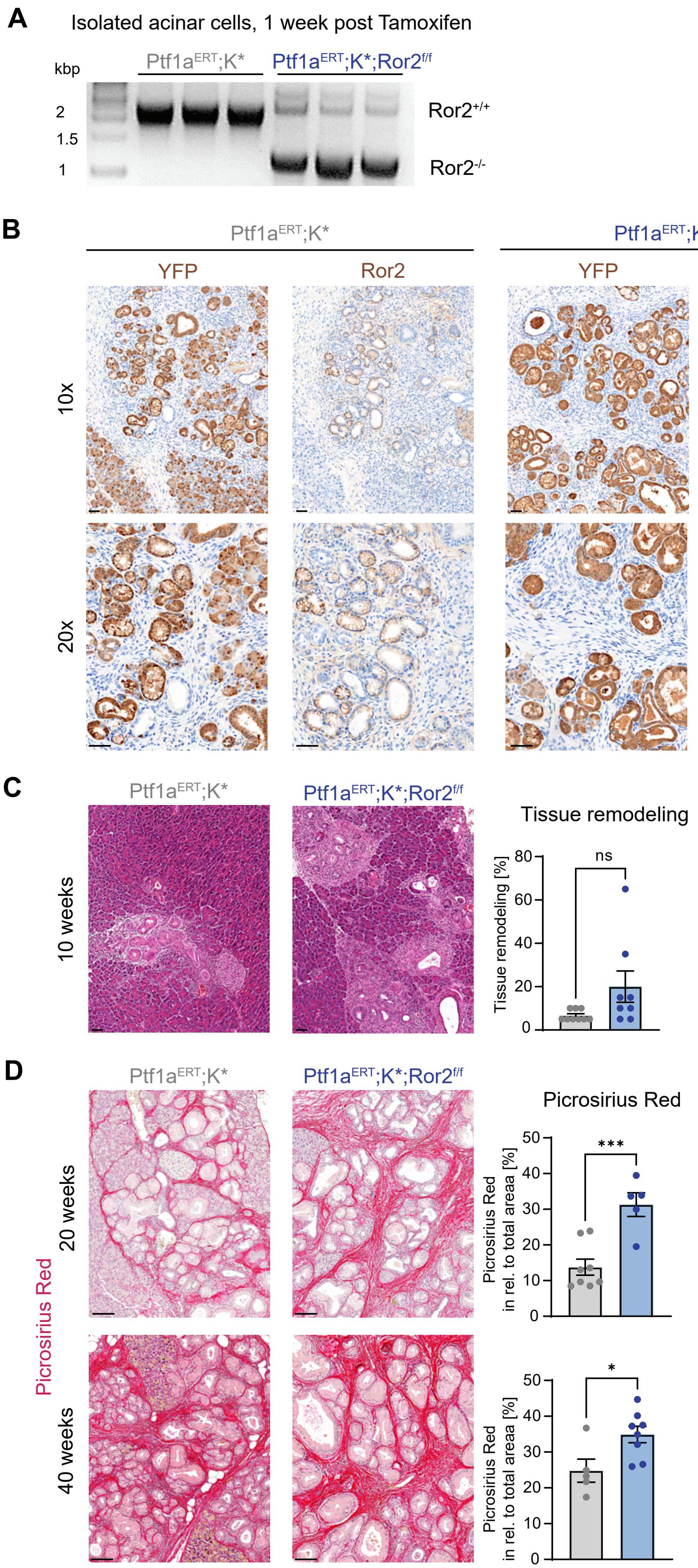

#### Supplementary Figure 5

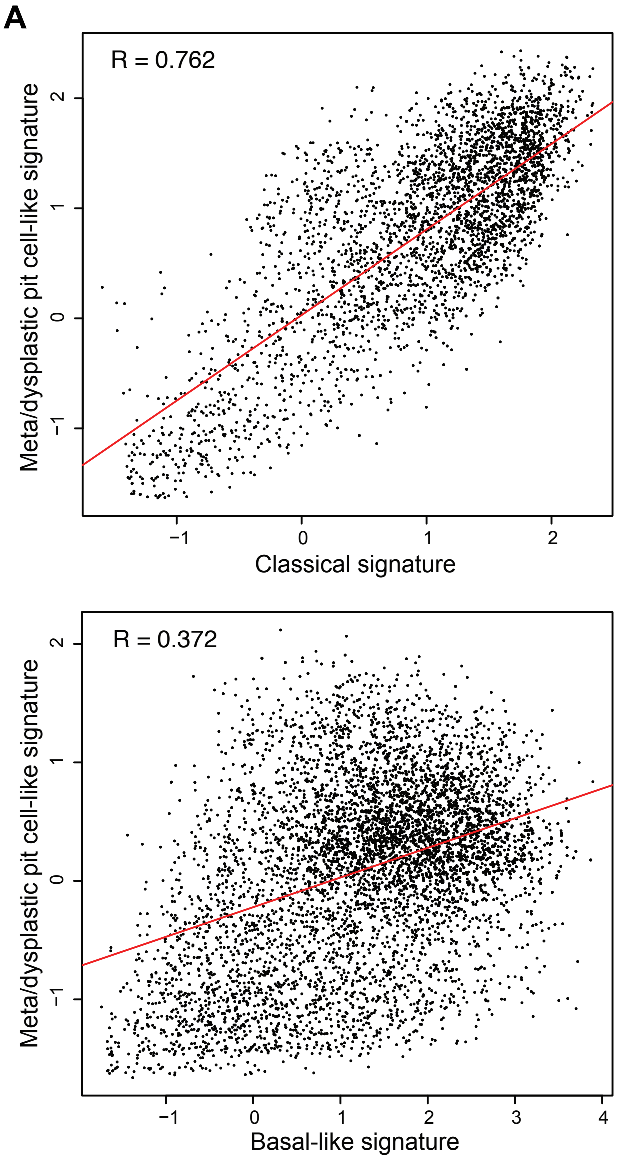


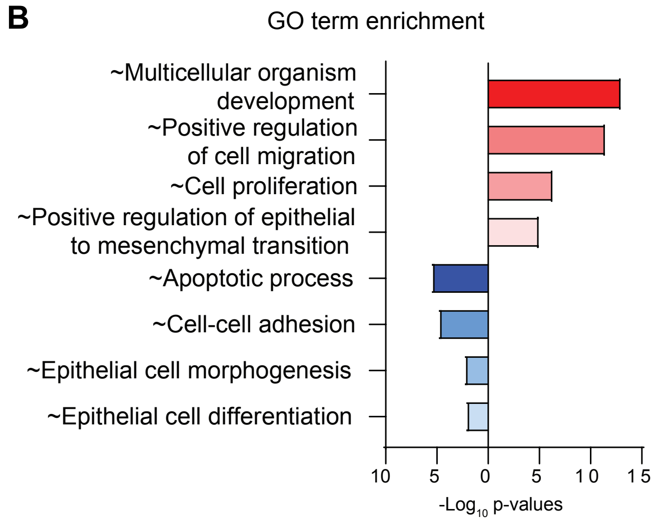

#### Supplementary Figure 6

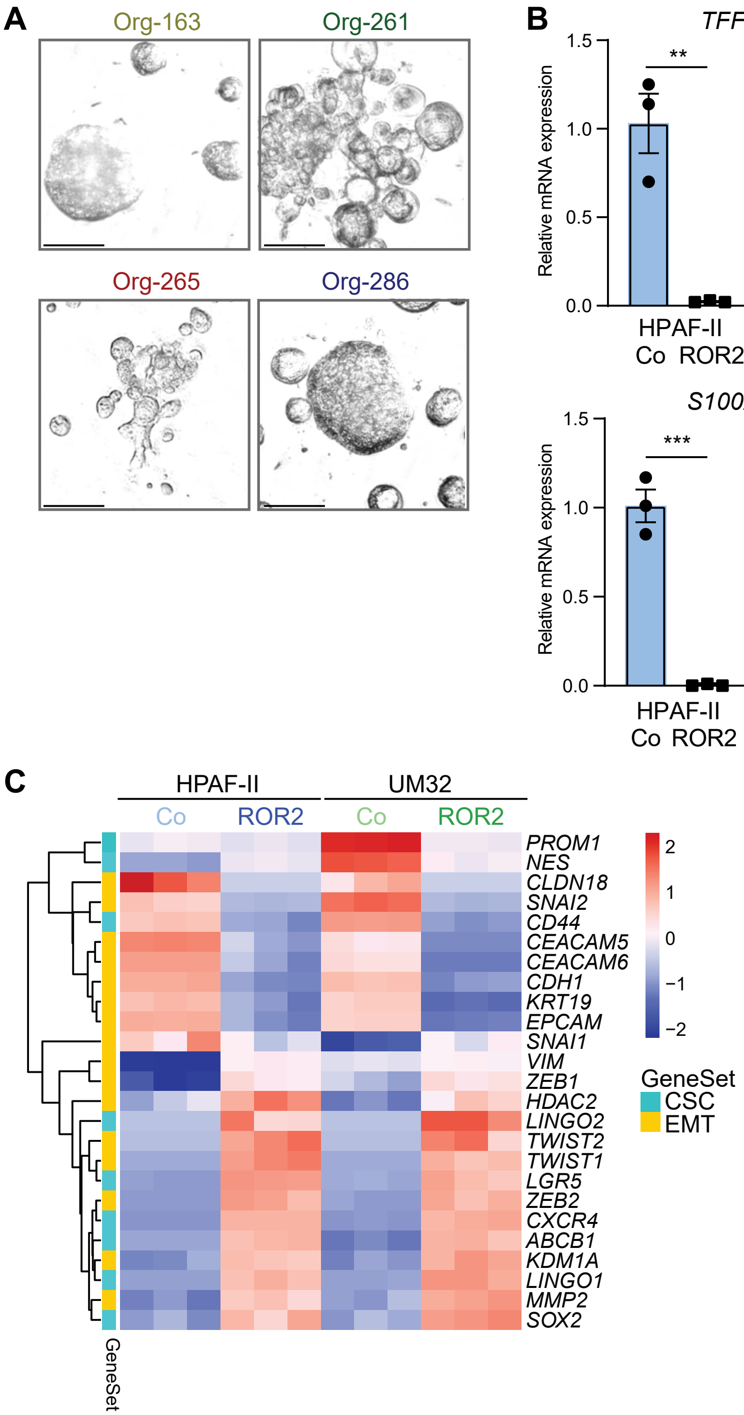


C

#### Supplementary Figure 7

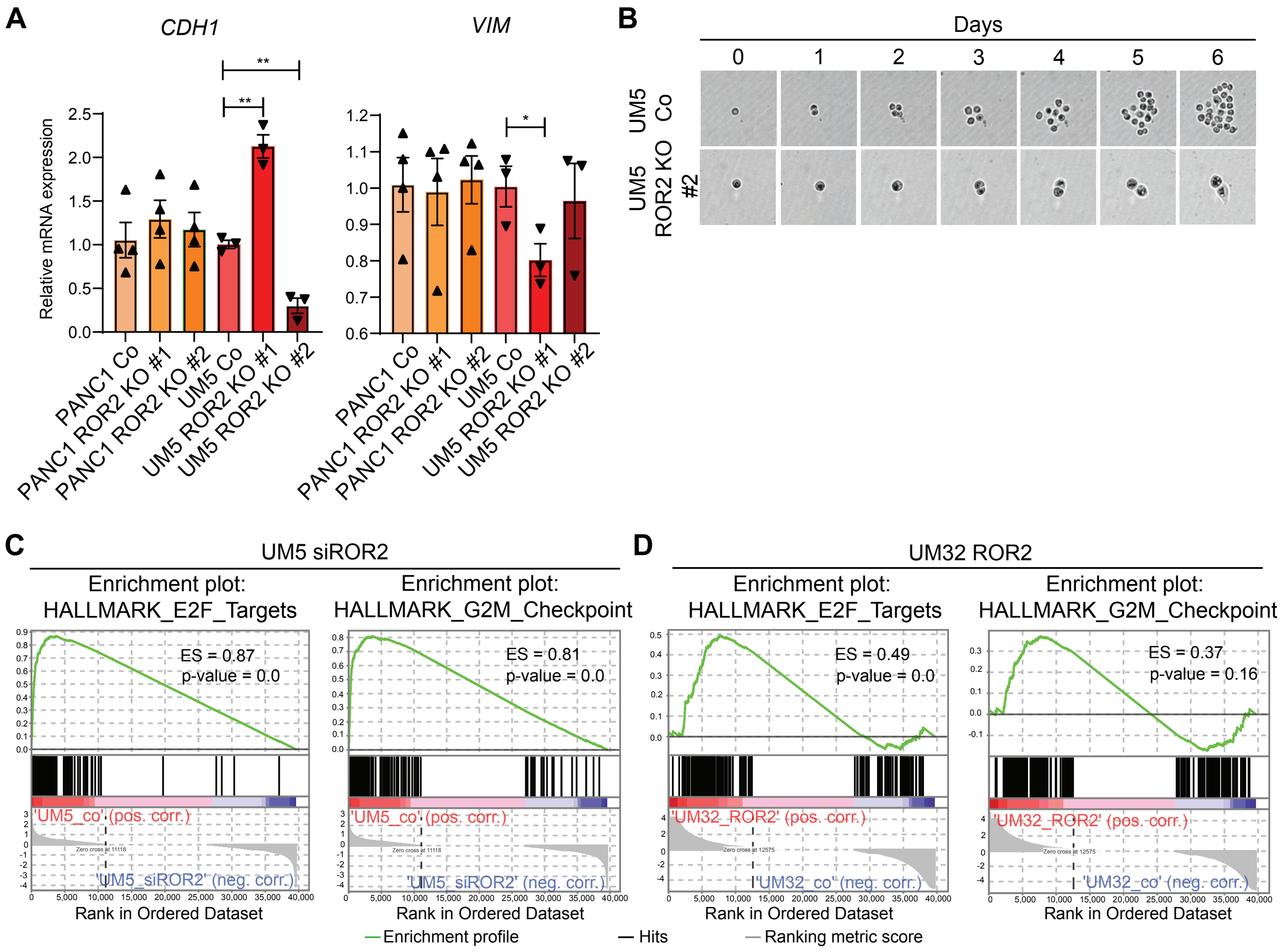

**Suppl. Table 1: Antibodies**

| Antibody | Application, dilution | Manufacturer, order number |
|---|---|---|
| AKT | WB: 1:1000 | Cell Signaling, 4691 |
| Aquaporin5 | IHC: 1:100 | Millipore Sigma, HPA065008 |
| α-Amylase | IHC: 1:800 | Sigma-Aldrich, A8273 |
| E-Cadherin | WB: 1:2000 | Cell Signaling, 3195 |
| E-Cadherin | IF: 1:200 | BD, 610182 |
| ERK | WB: 1:1000 | Cell Signaling, 9102 |
| GAPDH | WB: 1:2500 | Cell Signaling, 2118 |
| GFP/YFP | IHC: 1:200 | Cell Signaling, 2956 |
| Ki67 | IHC: 1:1000 | Abcam, ab15580 |
| KRT19 | IF: 1:200<br>IHC: 1:4000<br>WB: 1:5000 | Abcam, ab133496 |
| MUC5AC | IHC: 1:1000 | Fisher Scientific, MS145P1 |
| Phospho-AKT | WB: 1:1000 | Cell Signaling, 4060 |
| Phospho-ERK | WB: 1:1000 | Cell Signaling, 9101 |
| ROR2 | WB: 1:800<br>IF/IHC: 1:50 | Cell Signaling, 88639 |
| CDKN2A | IHC: 1:100 | Abcam, ab211542 |
| TFF1 | IHC: 1:100 | Abcam, ab190942 |
| Vimentin | WB: 1:2000<br>IF: 1:200 | Cell Signaling, 5741 |
| Vinculin | WB: 1:2500 | Cell Signaling, 13901 |
| ZEB1 | WB: 1:1000 | Abcam, ab203829 |

**Suppl. Table 2: Primer sequences**

| Target | Forward | Reverse |
|---|---|---|
| *Cela1* | GTGGACAGCCCCATGACTTA | TCATAGCCTGCAACCACGTT |
| *Junb* | AGGCAGCTACTTTTCGGGTC | TTGCTGTTGGGGACGATCAA |
| *Pdx1* | AAATCCACCAAAGCTCACGC | GGTCAAGTTCAACATCACTGCC |
| *Ptf1a* | CTTGCAGGGCACTCTCTTTC | CGATGTGAGCTGTCTCAGGA |
| *Ror2* | CGTGGTGCTTTACGCAGAAT | GCCATCTCGGGGACTACAC |
| *CDH1* | ATTGATGAAAATCTGAAAGCGGCT | GTCTTTGTCTGACTCTGAGGAGTT |
| *GAPDH* | CTCATTTCCTGGTATGACAACGAA | GCAGTGAGGGTCTCTCTCTT |
| *LGALS4* | GCAGCGAGGAGAGGAAGAG | TTTCCATTTACCACCACCTTGT |
| *ROR2* | GTCCCGGACTTCAGGTGAAG | GTGATATTGTTTACTGGCTCCAGAA |

| | | |
|---|---|---|
| *S100A6* | CTCCCTACCGCTCCAAGC | GCCGGAGTACTTGTGGAAGA |
| *TFF1* | TCCCTCCAGAAGAGGAGTGT | CCGAGCTCTGGGACTAATCA |
| *VIM* | CGGGAGAAATTGCAGGAGGA | AAGGTCAAGACGTGCCAGAG |
| *ZEB1* | GAACACACAGGTAAAAGACCTCAT | CACATTTGTCACATTGATAGGGCT |
